# Drug combinations targeting FAK and MEK overcomes tumour heterogeneity in glioblastoma

**DOI:** 10.1101/2024.11.26.625442

**Authors:** Muhammad Furqan, Richard J R Elliott, Peter Nagle, John C. Dawson, Roza Masalmeh, Virginia Alvarez Garcia, Alison F. Munro, Camilla Drake, Gillian M Morrison, Steven M. Pollard, Daniel Ebner, Valerie G. Brunton, Margaret C Frame, Neil O. Carragher

## Abstract

Glioblastoma (GBM) is an aggressive brain tumour with limited treatment options and poor prognosis, largely due to its heterogeneity and the involvement of multiple intracellular signalling pathways that contribute to drug resistance. Standard therapies have not significantly improved patient outcomes over the past two decades. While recent advancements in targeted drug combination therapies, such as dabrafenib and trametinib, show promise for certain GBM subgroups, identifying drug combinations effective across the broader GBM population remains a challenge. Integrin-mediated signalling, particularly through Focal Adhesion Kinase (FAK), plays a pivotal role in GBM pathogenesis and invasion, making it a potential therapeutic target [1].

In our study, we utilized a chemogenomic screening approach to identify synergistic drug combinations that target FAK in glioblastoma. We initially employed a CRISPR-engineered GBM model to assess the effects of FAK depletion and discovered that combining FAK inhibitors with MEK inhibitors, particularly trametinib, demonstrated synergistic effects. This potent combination was validated through various 2D & 3D assays, including cell viability/apoptotic assessment, synergistic analysis, cellular imaging, and target engagement assays. The combination also effectively inhibited spheroid growth and invasion across a diverse panel of patient derived GBM stem cells. Molecular mechanisms underlying these effects included suppression of multiple kinase signalling pathways and enhanced apoptosis, elucidated using Reverse Phase Protein Array (RPPA) profiling and western blot validation. *In vivo*, the combination therapy significantly reduced tumour volume in orthotopic transplantation models. These findings suggest that combining FAK and MEK inhibitors represent a promising therapeutic strategy to overcome the challenges of GBM treatment.

## Introduction

Glioblastoma (GBM) remains a cancer of high unmet clinical need despite considerable efforts to develop new treatments. The current standard of care for newly diagnosed GBM patients consists of surgery, temozolomide (TMZ), and ionizing radiation (IR) which provides an overall 15-month survival benefit and has not been improved upon since the EORTC-NCIC trial published in 2005 [2]. The self-renewing capacity of GBM stem cells is an inherent feature of GBM to which the tumour microenvironment contributes supporting adaptive signal rewiring, heterogeneous tumour evolution and therapeutic resistance [3, 4]. Current treatment and clinical trials of modern targeted therapies have been largely ineffective because they fail to address inherent tumour heterogeneity contributing to incomplete treatment response and resistance due to the emergence of hypermutator phenotypes and rapid rewiring across multiple signalling pathways [5, 6].

Large-scale genomic analyses performed over the past decade has enhanced our understanding of GBM biology supporting classification of GBM into different subtypes based on gene signatures that may help predict prognosis and therapeutic response [7-9]. However, this new understanding of the GBM genomic landscape has not yet yielded effective personalized medicine strategies across the GBM population predominantly as a result of the fact that GBM is driven by multiple intracellular signalling pathways rather than single gene alterations. Thus the challenge in GBM is identifying effective drug combinations which overcome tumour heterogeneity and therapeutic resistance by targeting the tumour’s ability to re-wire its signalling networks following treatment with single pathway blocking therapy [5].

Despite the challenges, new hope is provided from the recent Food and Drug Administration (FDA) approval of the combination of the targeted BRAF inhibitor dabrafenib and MEK inhibitor trametinib for the treatment of nearly any type of advanced solid tumour that has a BRAF^V600E^ mutation [10]. Studies have shown that treatment of tumours with BRAF inhibitors (such as dabrafenib) develop resistance by activating signals from the MEK protein [11]. Thus, combining dabrafenib with the MEK inhibitor trametinib was postulated to prevent tumours from using this escape mechanism and has demonstrated increased efficacy in preclinical tumour models and clinical studies [11-13]. The phase II Rare Oncology Agnostic Research (ROAR) clinical trial of dabrafenib plus trametinib in patients with BRAF V600E mutation-positive recurrent or refractory High Grade Glioma (HGG, including GBM), and Low Grade Glioma (LGG) constitutes part of a larger basket trial BRF117019 (NCT02034110), evaluating dabrafenib and trametinib combination therapy across multiple BRAF V600E-mutant tumours. The ROAR clinical trial primary endpoint of investigator-assessed overall response rate was 33% in the HGG cohort. Secondary endpoints of median duration of response was 31.2%, progression-free survival, 5.5% and median overall survival was 17.6 % in HGG [14]. While only 2% of adult GBM express BRAF V600E the incidence of activating oncogenic BRAFV600E mutation is higher across other central nervous system primary tumours in paediatric and adult patients [15], thus dabrafenib combined with trametinib represents a potential alternative treatment option for these subsets of patients.

With an almost infinite multi-drug possibilities, identifying the most effective drug combinations across other GBM patient subgroups presents a significant challenge as evidenced by the failure of the majority of drug combination trials performed in GBM to date [16]. The scientific rationale and preclinical evidence supporting many drug combination trials are often unclear and it appears that many drug combination strategies are opportunistic with selection based on pragmatic considerations including drug access and often representing an after-thought to rescue poor response of single agent therapy. Recent high throughput systematic testing of drug combinations across cancer cell models indicates that synergy between drugs is rare and highly context-dependent [17]. Thus, application of more evidence-based approaches to the discovery of novel drug combination strategies incorporating GBM models that represent the heterogeneity of disease are required to validate and prioritize the most promising drug combination strategies for clinical development.

Altered extracellular matrix (ECM) remodelling and intracellular signalling mediated by the integrin family of ECM receptors and their downstream kinases effectors such as Focal Adhesion Kinase (FAK) has been proposed as an important mechanism influencing GBM pathogenesis, formation of the tumour niche and brain tissue invasion [18]. Overexpression of integrin receptor subunits, β1, β3, α3, α5, αv and high expression of FAK and its closely related homolog PYK2 are associated with reduced overall survival of GBM patients [19-22]. However, therapeutic strategies to directly inhibit aberrant integrin-mediated ECM adhesion and signalling, as exemplified by the peptidomimetic drug candidate cilengitide, which blocks the ECM-RGD binding site of αvβ3 and αvβ5 integrins failed to improve overall survival of patients with GBM despite promising preclinical data [23, 24]. The failure of such integrin-targeting therapy has been proposed to be a consequence of a variety of pro-survival bypass signalling pathways mediated by multiple other integrin receptors [18] and other upstream ECM adhesion signalling receptors known to be dysregulated in GBM including discoidin domain receptors (DDR) [25] and CD44 [26]. These findings suggest that targeting downstream mediators positioned at the intersection of multiple integrins and receptor tyrosine kinases signalling pathways such as FAK may have additional therapeutic benefit. We and others have demonstrated that FAK is an important downstream mediator of invasion and survival pathways in multiple cancer cell types including GBM [27-29]. In human GBM cell models both small molecule pharmacological inhibition and genetic knockdown of FAK suppress the proliferation, survival, and 3D invasion of glioma stem cells [30, 31]. Pharmacological inhibition of FAK has also been shown to enhance the effect of temozolomide on tumour growth in a C57BL/6-GL261 mouse glioma model [32] and displays radiosensitizing effects in a subset of GBM cell models [30]. Furthermore, in a murine *in vivo* surgical resection model FAK and PYK2 signalling are upregulated in recurrent tumour samples [22]. Treatment with the small molecule PYK2/FAK inhibitor PF-562271 reversed PYK2/FAK signalling activation in recurrent tumours, reducing tumour volume by 43%, and increased animal survival by 33% [22]. While these results are promising, more effective, and durable responses would be desirable. Collectively, these studies indicate that FAK is a promising therapeutic target in GBM but may only yield maximum patient benefit when exploited as part of a synergistic therapeutic drug combination.

In this study we employed an unbiased chemogenomic screening approach in an isogenic model of FAK-kinase deficient (KD) and FAK-wild-type (WT) expressing GBM cells to identify drug combinations that could potentially synergise with depletion of FAK kinase activity. Our screen identified a number of drug candidates including four distinct MEK inhibitors that enhance GBM cell death when combined with FAK-KD cells, relative to FAK-WT. Focussing on the most clinically advanced MEK inhibitor trametinib, we demonstrate low level synergistic activity upon FAK-WT GBM cell survival and invasion endpoints when combined with multiple small-molecule FAK inhibitors. Post-translational pathway profiling indicates that trametinib promotes multiple compensatory signalling and pathway rewiring events in GBM cells when used as single agent. Validation studies across a heterogenous panel of patient-derived GBM stem cells demonstrate potent FAK+MEK inhibitor combination activity at nanomolar concentration across 2D and 3D phenotypic assays and reduced tumour area in an *in vivo* orthotopic transplantation model. Our systematic GBM phenotypic discovery platform demonstrates the robustness of FAK+MEK inhibitor combination treatment across heterogenous GBM stem cell models, supporting the case for further preclinical development toward clinical studies.

## Material and Methods

### Cell culture

To explore the role of FAK in GBM, we utilised a recently-described transformed neural stem cell-derived GBM cell model featuring CRISPR/Cas9-mediated deletion of the tumour suppressor genes *Nf1* (N) and *Pten* (P), and concomitant expression of the constitutively active, oncogenic EGFR (E) variant EGFRVIII (hereafter termed NPE cells)[33]. To investigate the function of FAK in NPE cells, we performed CRISPR/Cas9-mediated knockout of FAK in NPE cells and re-expressed FAK^K454R^ kinase deficient mutant protein (NPE-FAK-Kinase deficient or -KD)) and wild-type FAK protein at endogenous levels (NPE-FAK-WT) in the same clonal population (see Supplementary Figure 1), thus creating paired isogenic NPE cells whose phenotypic differences can be attributed solely to expression catalytically active FAK.

Patient derived glioblastoma stem cell lines used in this study were obtained from the Glioma Cellular Genetics Resource, Edinburgh (supplementary table S1). Cells were maintained in DMEM/HAMS-F12 media (Sigma, D8437) supplemented with glucose (8 mM, Sigma G8644), beta-mercaptoethanol (100 uM, Gibco, 31350-010), MEM non-essential amino acids (Gibco,11140-035), BSA (0.015%, Gibco 15260-037), recombinant hFGF basic (12 ng/mL, Peprotech 100-18b), recombinant mEGF (12 ng/mL, Peprotech, 315-09) and laminin (4-10 ug/mL, R&D Stems 3446-0050-01), B27 (1:100, Gibco 17504-044) and N2 supplements (1:200, Gibco 17502-048) [34]. Cells were plated onto 384-well microplates (Greiner Bio-One, 781091), after pre-coating with laminin (10 µg/mL in media), at 500-1500 cells/well.

### Compound and drug combination screening

LOPAC Library (1280 compounds, Sigma Aldrich #LO4100) plates were supplied as assay ready microplates (10mM stock in DMSO, 1.5ul per well), by BioAscent Ltd. The custom C3L library [35] was supplied direct via our partners at the University of Oxford (Target Discovery Institute). Compound plates were diluted in media (1:50 dilution, 75uL/well, 200uM, 2% DMSO (v/v)) using a ViaFill liquid dispenser. The plates were transferred to a BioMek Liquid handling platform and serially diluted to 20X the final concentrations required (see below, in media containing 2% DMSO (v/v)). Staurosporine (1uM) was added to column 1 of each compound plate as positive control for cell death. Finally, 2.5uL of each diluted compound was transferred to the cell plates (1: 20 dilution, final volume 50uL, 0.1% DMSO (v/v) in all wells including ‘untreated controls’ n=48) at final compound concentrations 1000, 300, 100, 30nM. The plates were incubated at 37°C and 5% CO2 for 48 hrs prior to viability assay.

Hit compounds were validated in 384w in triplicate by 8-point dose response (1000 to 10nM). Dose response profiles and IC50 ranges were used to build drug combination dose ratio matrices (384w format, 7x7 matrix per combination pair), using a D300e digital dispenser (Tecan.com). After 48 hrs, viability was quantified (PrestoBlue) and the data normalised to DMSO (% Cell Survival).

### Cell viability assay

NPE-FAK-WT and FAK-KD cells were seeded in 384-well plates at 400 cells per well and incubated for 24 hours before treatment. Cells were incubated with compounds for 48 hours; untreated cells were incubated with 0.1% DMSO as vehicle control (n= 48 per plate). PrestoBlue HS Cell viability reagent (5uL, Invitrogen; #P50200) was added to each well using ViaFill dispenser and plates incubated for 2 hours at 37°C. Fluorescence emission was read on an EnVision 2101 multilabel plate reader (PerkinElmer; ex540 nm, em590 nm). Background fluorescence was subtracted from ‘cell free’ wells. All conditions were then normalized on a plate-by-plate basis to internal DMSO controls (average of n = 48) to give % survival and Z-score quantification to compare relative survival (NPE-FAK-KD vs NPE-FAK-WT) and induced drug sensitivity in a FAK suppressed background.

### Cellular imaging and Quantification

Human glioma stem cells (GCGR cell lines) in 384w format were quantified at assay end points by nuclei staining and imaging using an ImageXpress high content microscope (Molecular Devices). Cells were fixed *in situ* by the addition of 8% formaldehyde (equivalent volume, cooled plates) using a multidrop dispenser (Thermo), incubated for 30 minutes and fixative removed using BioTek multi-well plate washer. The cells were then stained in Hoechst-33342 (H1399, Molecular Probes, 2ug/mL in PBS, 0.1% Triton-X100 (v/v)) at RT for 20 minutes, followed by washing and sealing with foil. Processed plates were then loaded into an ImageXpress-XL microscope using a KX robotic arm (PAA) and imaged with a 20X objective (6 sites per well). Total Nuclei counts were quantified using MetaXpress built-in analysis module (‘Count Nuclei’) (Molecular Devices). For representative images (Figure 4 & 5), fixed cells were stained with Hoechst (1:5000, 2ug/mL), Phalloidin-594 (ab176757, Abcam, 1:1500), Concanvalin A (C11252, Invitrogen, 30ug/mL). Images were acquired using an ImageXpress microscope, 20X objective.

### Quantification of tumour area in IHC stained sections

Tumour areas in immunohistochemically stained sections were quantified using ImageJ software. High resolution images of stained tissues were first converted to grayscale. Thresholds were adjusted to isolate positive staining, distinguishing it from the background. The software’s measurement tools were then used to calculate the area of the stained tumour regions, which was expressed as the total area of stained tissue section. This approach allowed for a consistent and objective quantification of tumour burden across different samples.

### Synergy calculation

Experiments were carried out in 384w format with multiple 7x7 dose combination matrices (including ‘drug alone’ arms). Typical dose ranges for the compounds used are VS4718, AMP-945 (500-100nM in 2D & from 3uM in 3D) and trametinib, GDC0623 (25-1nM, 2D, from 250nM in 3D). At assay end points (72 hrs for human glioma cells, 48 hrs for NPE derived lines), cells were fixed and stained with Hoechst (as above). Image quantification of nuclei counts, followed by DMSO normalization, gave % cell survival measurements which were then analyzed using SynergyFinder software3[36], incorporating multiple synergy models including Bliss, Loewe, HSA and ZIP synergy models for robust synergistic ranking with a ‘Combined Sensitivity Score’ (CSS)[37-39].

### 3D spheroid invasion assay

Cells were seeded in 96-well ultra-low-attachment “U” bottom plates (Corning; #7007) at 2000 cells per well in normal growth medium and centrifuged at 200 × g. Cells were allowed to form a single spheroid per well over 3-4 days before being overlaid with Matrigel (4 mg/ml). Subsequently, they were treated with drugs with continuous monitoring of spheroid outgrowth every 3 hrs over 12 days using an IncuCyte S3 live cell imaging system and analyzed with the built-in 3D Spheroid Invasion module. The imaging channels ‘Phase + Brightfield’ with a 10× objective were utilized to observe spheroid size and invasion

### Western blotting

Cells or spheroids were washed with PBS and lysed in ice cold RIPA buffer (50 mM Tris-HCl pH 8, 0.1% sodium dodecyl sulphate, 0.5% sodium deoxycholate, 1% Triton X-100, 150 mM NaCl) added with PhosSTOP™ Phosphatase Inhibitor (Roche; #4906845001) and cOmplete™ ULTRA Protease Inhibitor (Roche; #5892953001) cocktail for half an hour on ice with gentle agitation. Lysates were centrifuged at high speed to clear the debris; supernatants were quantified using Bradford assay reagent followed by protein normalization. Then, samples were resolved in 4%–15% Mini-PROTEAN TGXTM gels, transferred on to PVDF membrane (BIO-RAD; #1704156) and blocked in Roche block followed by incubation with primary antibodies overnight. Next day, membranes were washed with PBS-T and incubated with HRP linked secondary antibodies for 1 hour. Then, membranes were again washed three times with PBS-T and visualized using Clarity Western ECL substrate (BioRad; #1705061) on ChemiDoc (BioRad) imaging system. Antibodies used in western blot are listed in supplementary table S2.

### Animal studies

All procedures involving mice adhered to protocols approved by the Home Office in the United Kingdom under project license PP7510272 held by V.G.B. and were approved by the University of Edinburgh Animal Welfare and Ethical Review Body (PL05-21). 10-week-old female CD-1/nude mice were intracranially implanted with NPE cells (2x10^4^) or G7 cells (2x10^5^) in 2 µL growth medium following isoflurane anaesthesia. Buprenorphine was given for analgesia during surgery, and carprofen was administered in water for the subsequent 48 hours. Tumour cells were precisely implanted at coordinates 0.6 mm anterior and 1.5 mm lateral to the bregma, reaching a depth of 2.5 mm. Tumour growth was monitored biweekly by injecting 150 mg/kg of luciferase, followed by imaging using the IVIS® Lumina S5 system. Mice bearing established tumours were randomized into different treatment groups. The treatment regimen involved administering VS4718 (75mg/kg) twice daily (7 days a week) and/or trametinib (0.5mg/kg) once daily (5 days a week) for total of two weeks *via* oral gavage. Both the drugs were prepared in 0.5% hydroxypropyl methylcellulose solution. At the end of treatment, mice were humanely sacrificed through cervical dislocation, brains were collected and fixed in formalin overnight, followed by processing for paraffin embedding. Subsequently, sections were cut and subjected to stain with H&E and anti-GFP (for NPE tumours) and human anti-Mitochondria antibody (for G7 tumours) using conventional techniques.

### Immunohistochemistry

Immunohistochemistry was performed using DAKO (Agilent Technologies, Santa Clara, CA, USA) reagents. Briefly, mice brains were cut into two coronal sections near implantation sites followed by embedding in the paraffin blocks. 4uM thick brain slices were placed onto glass slides and dewaxed using xylene. Then, 10mM citrate buffer was used for antigen retrieval in a pressure cooker and slides were washed in water and TBS-T. Sections were then blocked with Dako REAL peroxidase block solution (Agilent; #S2023) and serum-free protein block solution (Agilent; #X0909) followed by overnight incubation in primary antibodies (anti-GFP 1:400) and (human anti-Mitochondria 1:100) (Novus, #NBP2-32980). Next day, sections were washed with TBS-T and incubated with DAKO EnVision-HRP rabbit/mouse (Agilent; #K5007) for 2 hours. After incubation, sections were again washed with TBS-T and visualized with DAB reagent (Agilent; #K3468). Finally, sections were counterstained with hematoxylin (Agilent; #S3309), dehydrated using ethanol and mounted using DPX mountant (Sigma-Aldrich; #06522).

### Reverse Phase protein Array

The Reverse Protein Array analysis was performed at the Cancer Research UK Scotland Centre’s Host and Tumour Profiling Unit. Cell lysates were mixed with 4× SDS sample buffer without bromophenol blue and supplemented with 10% 2-β-mercaptoethanol, at a final concentration of 2 mg/ml and final volume of 150 μl. Samples were stored at −80 °C prior to further analysis. Samples were denatured by heating to 95 °C for 5 min prior to printing as serial dilutions (1.50 mg/ml, 0.75 mg/ml, 0.375 mg/ml and 0.1875 mg/ml) in arrays consisting of 36 × 12 spots at a 500 μm spot-to-spot distance on the Aushon 2470 Arrayer Platform (Quanterix, MA, USA) using 8 × 185 μM pins, with 2 deposition rounds per feature. Each sample dilution series was spotted on all arrays with 8 arrays per slide on single pad Supernova nitrocellulose slides (Grace BioLabs, OR, USA). Sample loading on the slides, for normalisation purposes, was determined with Fast-Green dye staining and scanning using the InnoScan 710 slide scanner (Innopsys, Carbonne, France) at 800 nm. The RPPA slides were then washed with deionized water for 4 × 15 min on a platform shaker, prior to a 15 min incubation with Antigen Retrieval Reagent (1× Reblot strong). The slides then underwent two more 5 min washing steps with deionized water. Subsequently, the RPPA slides were placed in a ProPlate chamber (Grace BioLabs, OR, USA) in fresh deionized water and washed with 1× PBS twice for 5 min. The slides were then incubated for 10 min in the Superblock T20 Blocking Buffer (Pierce, Thermo Fisher Scientific, MA, USA), prior to TBST washing twice for 5 min. The RPPA slides were then incubated for 60 min with the 120 primary antibodies all diluted at 1:250 in Superblock buffer prior to two more TBST washes (5 min each) and a second 10 min incubation with the Blocking buffer, as described above. The RPPA slides were then incubated for 30 min with the DyLight-800-labelled anti-species secondary antibody diluted at 1:2500 in Superblock, followed by two 5 min washes in TBST and a final rinse with deionized water. Non-specific background signal was determined for each slide by performing the primary antibody incubation step solely with Superblock, without antibodies, on 1 array followed by fluorescent tracer secondary antibody. The slides were dried for 10 min at RT prior to imaging with an InnoScan 710 slide scanner (Innopsys, Carbonne, France). Microarray images were analysed using the Mapix software (Innopsys, Carbonne, France). The spot diameter of the grid was set to 270 μm. Background intensity was determined for each spot individually and subtracted from the sample spot signal, thus generating a net signal for each spot. Relative quantification of each analyte was determined for each spot by measuring fluorescence intensity. The validity of the serial dilutions was ensured by generating a linear fit curve from the 4-point dilution series for all samples, on all arrays, using a flag system where an R2 value was generated and R2 > 0.9 values were deemed good. Relative fluorescence intensity (RFI) values corresponding to relative abundance of total and phosphorylated proteins across the samples set were calculated and normalised to total protein by calculating the ratio of Antibody RFI/Fast-Green RFI for all samples.

## Results

### Chemogenomic screening to identify synergistic combinations with loss of FAK kinase activity

We sought to first identify novel drug combinations which synergize specifically with loss of FAK in GBM by using a chemogenomic screening strategy. We obtained isogenic glioma stem cell models, derived from mouse neural stem cells engineered to express constitutively active EGFRvIII and CRIPSR mediated deletion of *Nf1* and *Pten [33]* (NPE cells). This line was further engineered by CRISPR to express FAK^K454R^ kinase deficient mutant protein (FAK-KD) and a FAK-re-expression control, representing FAK^wild-type^ protein (FAK-WT) (see methods). FAK-WT and FAK-KD lines were adapted to 384-well format with optimal seeding densities, using a cell viability endpoint. Four separate drug libraries were selected, specifically with a drug re-purposing context in mind, and included pre-clinical and/or FDA approved compounds: the TargetMol anticancer set (L2110, 330 compounds), Kinase Chemogenomic set (KCGS[40], 187 compounds), LOPAC (1280 compounds, Sigma Aldrich) and a custom chemogenomic library set (789 compounds, ‘Comprehensive anti-Cancer small Compound Library, or C^3^L)[35], with dosing across 4 concentrations (1000, 300, 100, 30nM). Library dilutions and additions were carried out using a Biomek FX liquid handling robot (Beckman Coulter) with compound exposure held at 48 hrs and consistent DMSO concentration (0.1% (v/v)), and cell viability quantified. Data was normalised to DMSO wells and converted to a z-score for direct FAK-WT vs FAK-KD comparison (X-Y scatter plots, Figure 1A).

**Figure 1.**
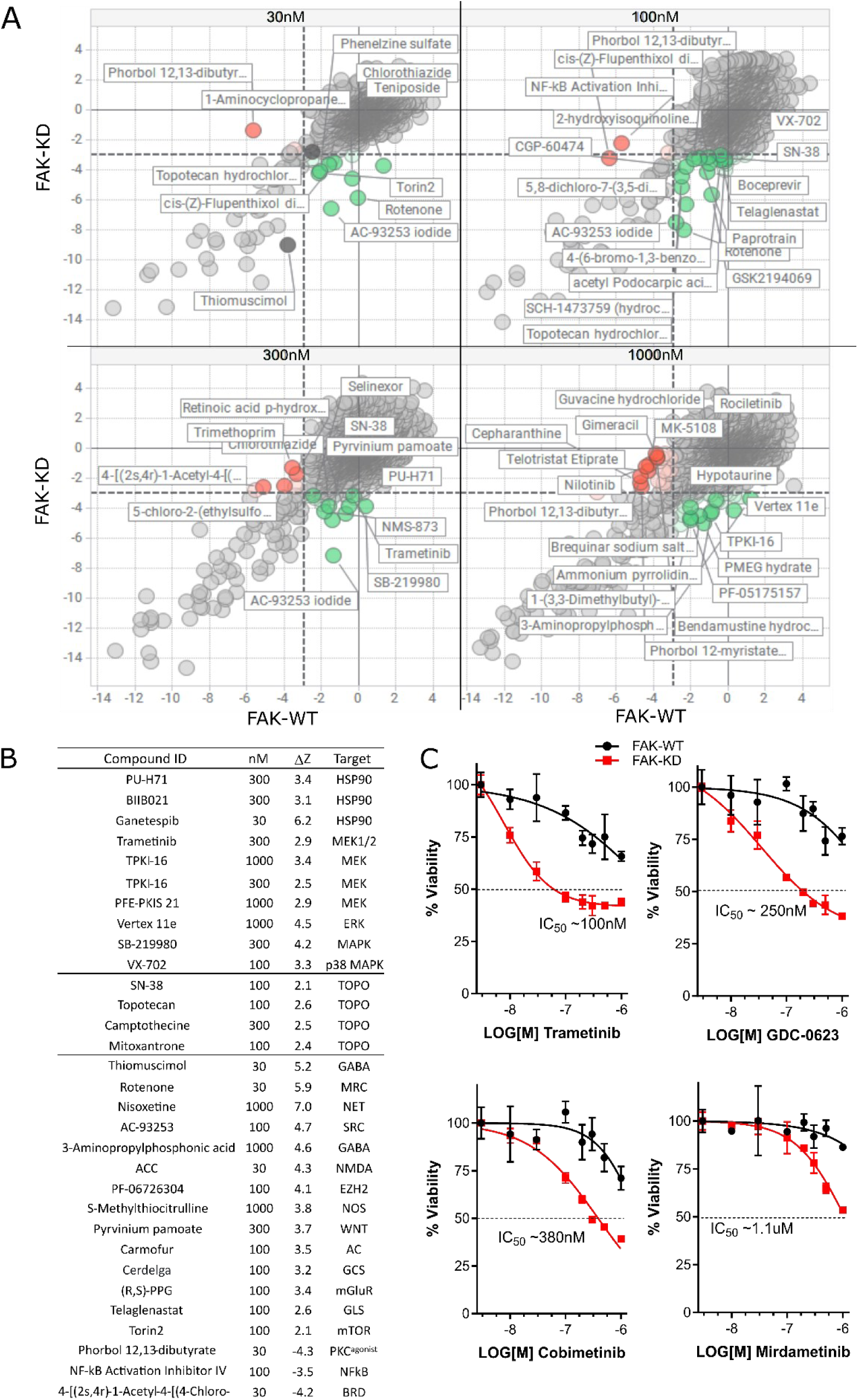
Chemogenomic screening against isogenic FAK-WT and FAK-KD cell lines. A. X-Y Scatter plots of normalised Z-scores of FAK-WT vs FAK-KD at 30, 100, 300, 1000nM. Key compounds are highlighted: FAK-KD sensitising – green, FAK-KD antagonists – red. B. ΔZ score rank was generated from thresholds of cell survival of FAK-WT cells (z score >-3) versus increased death in FAK-KD cells (z score <-3). Compound hit list with ΔZ scores >2 and higher are listed with their annotated target, plus example antagonistic hits ΔZ <-3. C. Validation of 4 MEK inhibitors (trametinib, GDC-0623, cobimetinib, mirdametinib) showing loss of FAK kinase activity (red curve) sensitises cells to MEK inhibition.

Quality control metrics were assessed on a plate-by-plate basis, specifically, robust Z-prime values (based on DMSO vs staurosporine controls) and normal sample distributions (Supplementary Figure 2). Compounds which showed increased death in the presence of inactivated FAK (FAK-KD), relative to FAK-WT, were assessed by differential analysis, generating a delta-Z score (ΔZ = WT z-score – KD z-score). Hit compounds were identified and ranked by a ΔZ score >2 (Figure 1B) and complete ranking is presented in supplementary Table 3. Top ranked hits included previously reported HSP90 inhibitors [41] and a number of MEK inhibitors, including the clinically advanced drug, trametinib. A number of MEK inhibitors were repurchased and validated (Figure 1C), with trametinib (MEK1/2), GDC-0623 (MEK1), cobimetinib (MEK1), and mirdametinb (pan-MEK) selectively inducing cell death on an inactive FAK kinase background, relative to FAK-WT cells.

### Identifying synergistic combinations between MEK and FAK inhibitors

A number of FAK inhibitors are currently undergoing clinical development, but have demonstrated limited ant-cancer activity when used as single agents, indicating that FAK inhibitors will be most effective when used in combination with other agents [42]. In order to phenocopy kinase-deficient FAK, we used the well characterized FAK inhibitor VS4718 to determine if there is synergistic activity in FAK competent WT cells, when used in combination with the MEK inhibitors GDC0632 and trametinib. We dosed FAK-WT by 7x7 combination matrices. FAK-WT cells were prepared in both 2D and 3D (Matrigel supported) culture, followed by dosing and end point quantification of cell viability. Normalised data (% Cell survival) was analysed for synergy using SynergyFinder software (synergyfinder.org).

Drug combination data on NPE-FAK-WT cells indicated synergy/additivity when targeting MEK and FAK by chemical inhibition in both 2D and 3D models (Figure 2A-E), using either trametinib or GDC-0623.

**Figure 2.**
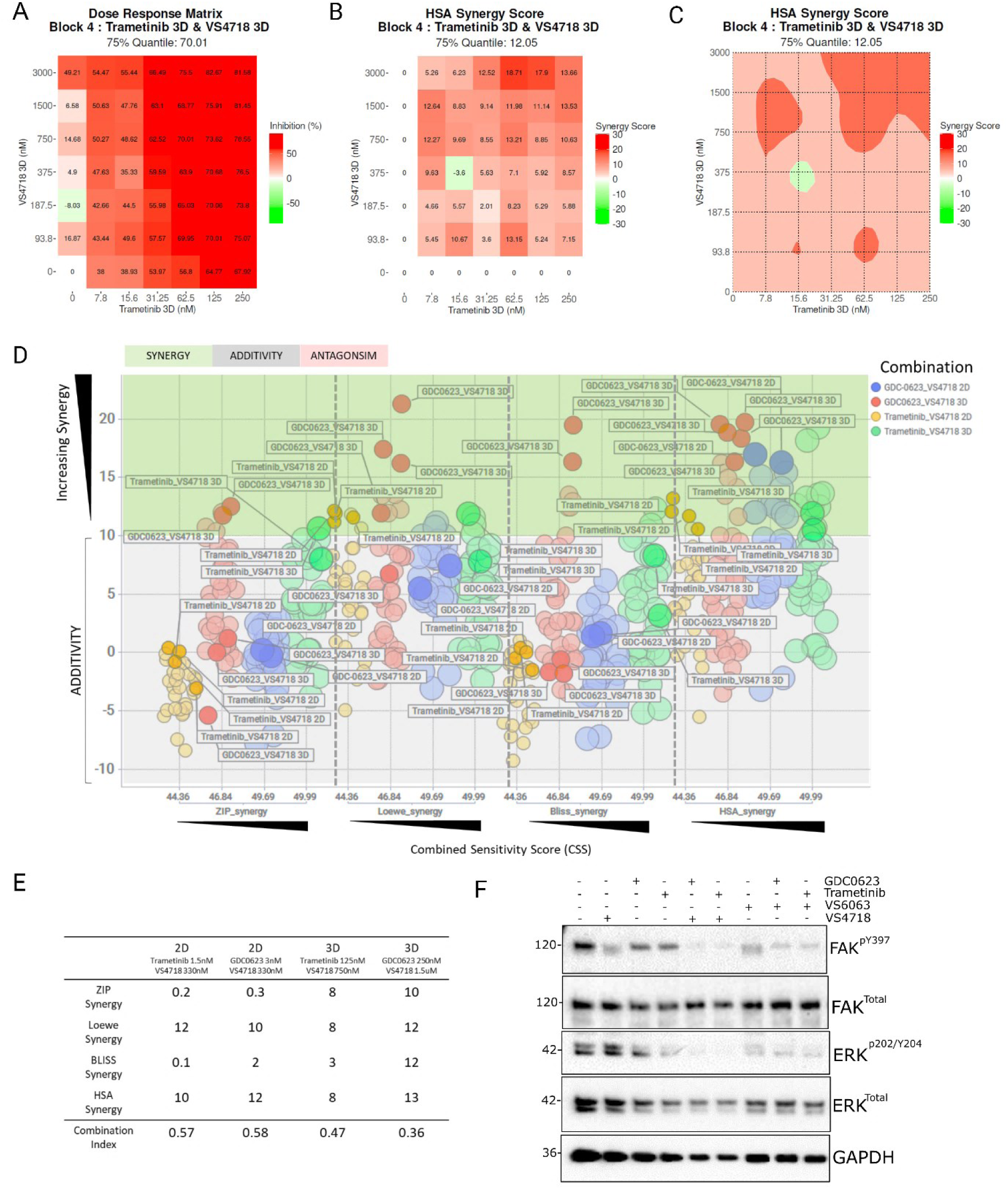
Chemical inhibition of MEK and FAK indicates additive-to-synergistic trends in 2D and 3D NPE-FAK-WT cells. A. Heatmap of % cell survival of combination matrices of VS4718 vs trametinib in 3D spheroids. B-C. Corresponding synergy heatmap and synergy contour plots (HSA model) of trametinib & VS4718 combination matrices, with increasing synergy indicated in red D. Scatter plot of Synergy (green zone)/additivity (grey zone) vs potency (CSS) across 4 synergy models (BLISS, ZIP. Loewe, HSA), from combinations on NPE-FAK-WT cells (2D & 3D) with trametinib or GDC0623 vs VS4718. F. Western blot of validation of on-target trametinib and VS4718 inhibition in this cell model.

A representative heatmap of the FAK+MEK inhibitor combination matrix showing % Inhibition (Figure 2A) and corresponding synergy score and contour map (Figure 2B, C) is shown, where synergy is indicated by maximal or peak synergy scores in darker red (Synergy scores ∼10+ across 4 synergy models preferred). Synergy scores across all combination doses in 2D and 3D are shown (Figure 2D), with three synergy models indicated (BLISS, ZIP, Loewe & HSA[36])[37, 38] and the synergistic zone shaded in green. The X-axis indicates Combination Synergy Score (CSS) which quantifies the potency of the combination (ie significant death caused by the synergy observed, indicated by a CSS >30) [39]. Trametinib combinations with VS4718 in 2D & 3D are indicated in yellow and green data points, GDC0623 plus VS4718 in 2D & 3D in blue and red data points. The most synergistic dose combinations which give combination index (CI) values <1 are shown in the figure 2E, with the responses in 3D showing greater effects across all 4 synergy models. Functional and on-target validation of MEK and FAK inhibitors used on this GBM cell model was carried out, with reduced phosphorylated (p)-FAK and phosphorylated (p)-ERK observed by Western Blotting (Figure 2F).

### FAK+MEK drug combination *in vivo*: investigation with mouse glioma model

To evaluate whether the combination of VS4718 with trametinib results in greater inhibition of tumour growth in mice, we utilized the NPE model which is known to form tumours within two weeks following intracranial implantation [33]. Initially, we confirmed drug target engagement by administering three different concentrations of trametinib (0.1, 0.5, and 1 mg/kg) alongside a previously established concentration of VS4718 (75 mg/kg). Both drugs were administered *via* oral gavage for three days. Western blot analysis of tumour lysates showed effective inhibition of p-FAK (Y397) at 75 mg/kg VS4718 and p-ERK (T202/Y204) at 0.5 mg/kg trametinib (Supplementary Figure 3).

To assess tumour growth inhibition, we treated NPE-FAK-WT tumour bearing mice with the individual drugs (VS4718 and trametinib), their combination, and a vehicle control (HPMC) for two weeks, followed by culling (Figure 3A). Immunohistochemical analysis of brain sections revealed weak responses to monotherapy, with significant tumour reduction observed only in the combination therapy group (Figure 3B). Quantification of tumour areas, based on tumour-specific GFP staining, showed a statistically significant reduction in tumour size with combination therapy compared to either monotherapy (Figure 3C). It is important to mention that the combination of higher doses of trametinib (1 mg/kg) with VS4718 (75 mg/kg) likely induced systemic toxicity, as indicated by significant weight loss in the treated mice (data not shown). In contrast, at the lower dose of trametinib (0.5 mg/kg) combined with VS4718 (75 mg/kg), only one out of five mice exhibited weight loss by the end of the experiment (Supplementary figure 4). Our results are consistent with findings from similar studies, where the combination of FAK and MEK inhibitors, despite their potent anticancer effects, has been associated with increased toxicity, particularly under more aggressive dosing regimens [43, 44].

**Figure 3.**
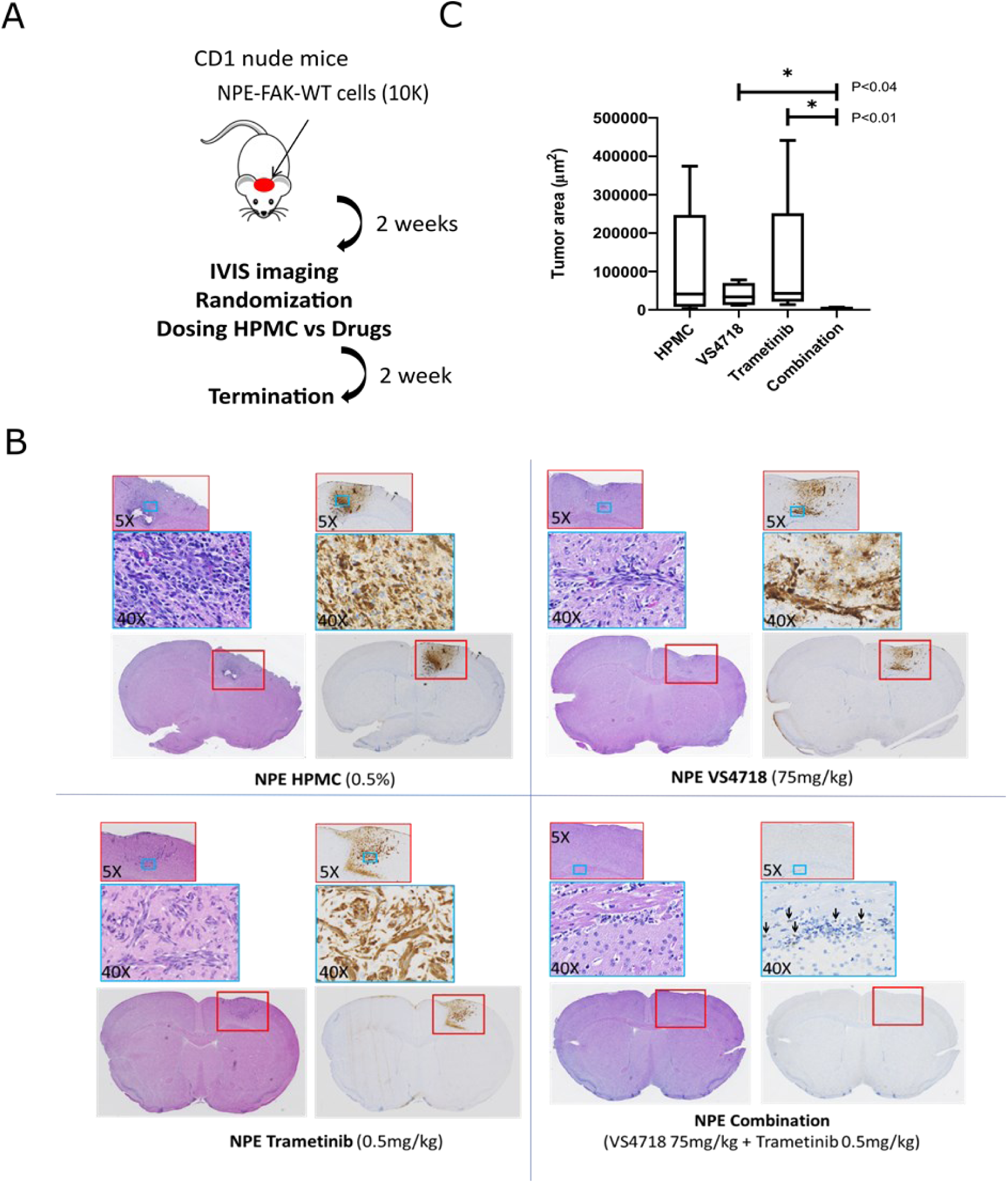
*In vivo* validation of FAK+MEK inhibitor synergy in NPE FAK-WT tumour bearing mice. **A** Schematic showing experimental protocol where mice with established tumours were treated with drugs (VS4718 & trametinib) or vehicle (HPMC). **B** Representative images of paraffin embedded brain sections containing NPE-FAK-WT tumours stained with Haematoxylin and eosin or following immunohistochemistry with anti-GFP antibody. Black arrows indicate GFP stained tumour cells in mouse brain. **C** Quantification of Tumour Area based on GFP staining.. n=5 each group. Error bars represent the standard error of the mean (SEM). * Indicates statistical significance at P < 0.0132.

### FAK & MEK Inhibitor profiling across patient derived human glioma stem cells

We further characterised MEK and FAK inhibitors across a panel of 14 human GBM stem cells (Figure 4). Individual dose response profiles of MEK inhibitors, trametinib (MEK1/2), GDC-0623 (MEK1), and FAK inhibitor VS4718, were carried out across the human glioma panel. Generally, less than 50% cell death was achieved at up to 1μM (VS4718) & 300nM (MEKi) after 72 hrs exposure (Figure 4A). Hierarchical clustering, demonstrating drug sensitivity profiles across the GBM stem cell panel for each compound vs normalised Area Under the Curve (AUC), is shown (Figure 4B).

**FIGURE 4:**
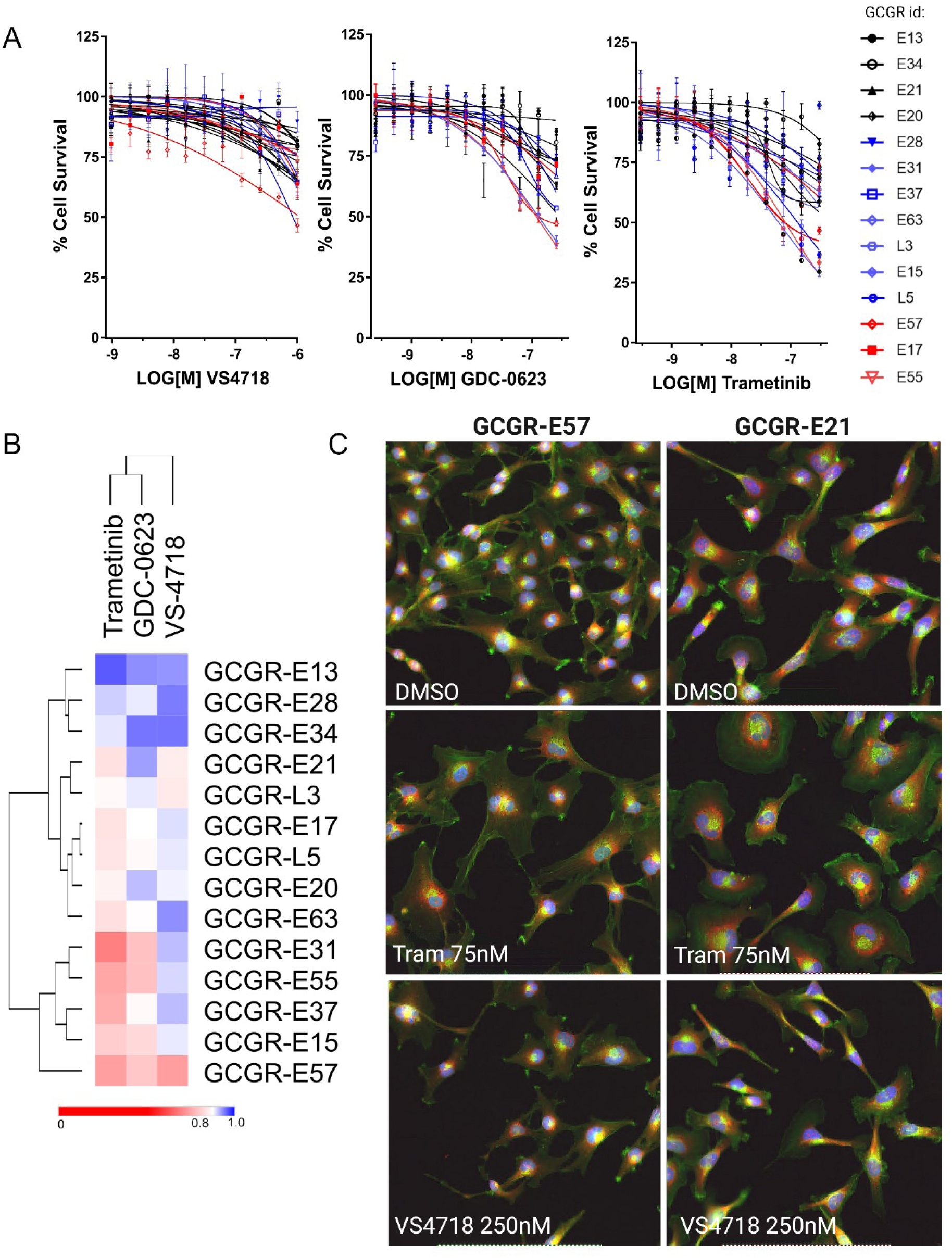
A. Dose response validation of FAK inhibitor (VS4718) and MEK inhibitors (trametinib & GDC-0623) across 14 patient derived glioma stem cell (GSC) lines. B. Heatmap and hierarchical clustering of dose response data by normalised Area Under the Curve (AUC). AUC data is scaled between 1 (100% inactive) and 0 (100% active). C. Representative images of E57 and E21 cells in response to VS4718 and trametinib at indicated doses (72 hrs). Staining by Hoechst (Nuclei, blue), Concanavalin A (ER, red), Phalloidin (f-actin, green). Scale bar indicates 20um.

Representative images of trametinib (75nM), VS4718 (250nM) phenotypes versus DMSO (0.1% (v/v)) are shown for GBM stem cells belonging to the mesenchymal subtype, GCGR-E21 and -E57 cells, indicating increased cell spreading upon trametinib treatment (Figure 4C). We further examined FAK and MEK inhibitor dose combinations across 14 GBM stem cell lines, as 7x7 dose combination matrices, in order to further investigate FAK+MEK inhibitor-synergy across a heterogeneous panel of human GBM stem cells (Figure 5).

**Figure 5.**
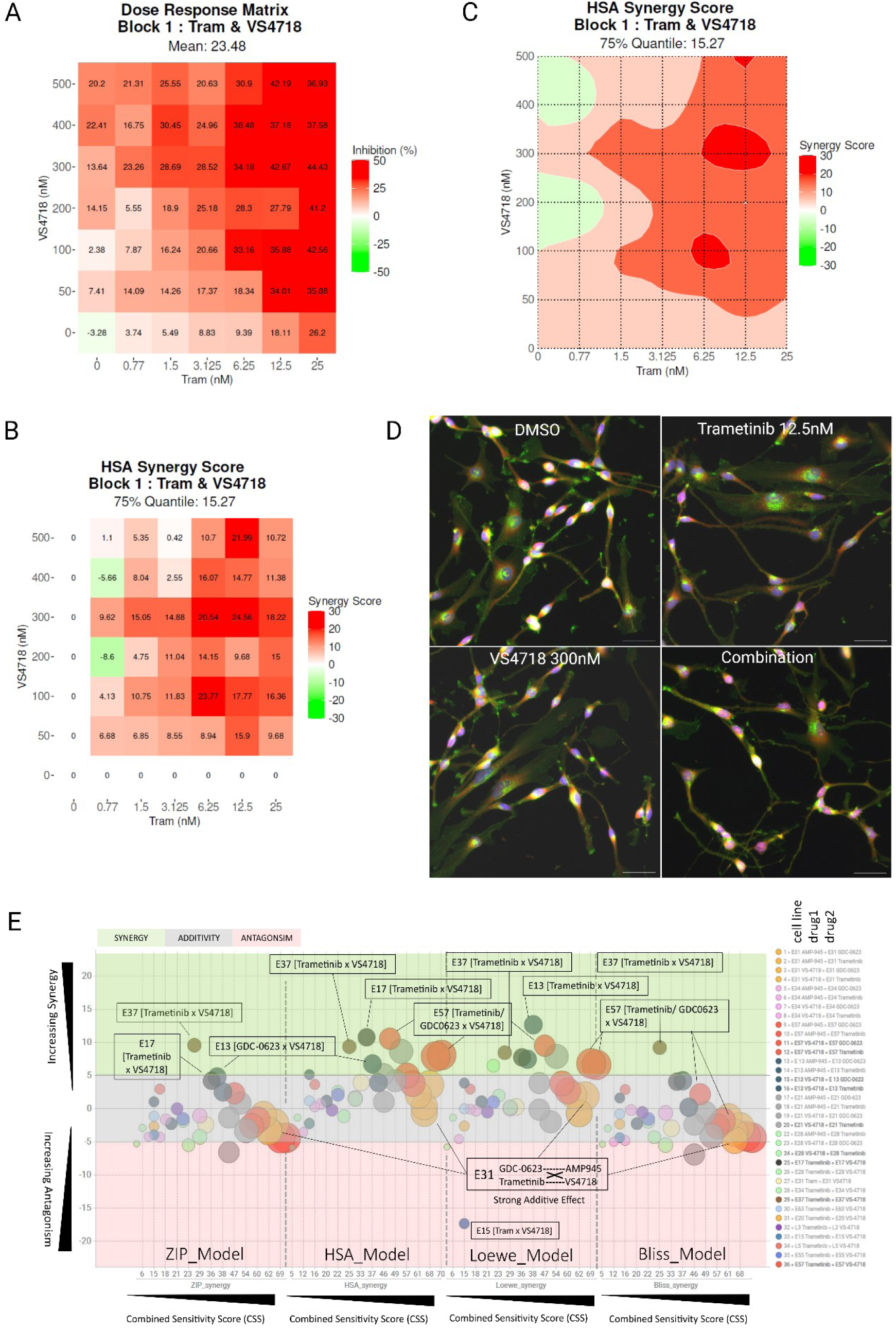
Combination screening (7x7 dose combination matrices) of MEK inhibitors (trametinib, GDC-0623) with FAK inhibitors (VS-4718, AMP-945) against a panel of 14 patient derived, human glioma stem cell lines. A. Representative data: dose combination matrices of % Inhibition (Max Red). B. Synergy combination plot (HSA Model) showing areas of max synergy (red) >10 δmax and C. Corresponding 2D contour plot showing ideal dose range (dark red zones). D Representative images of E17 cells treated with DMSO (0.1% v/v), trametinib (12.5nM) alone, VS-4718 (300nM) alone and in combination (synergistic doses) Staining by Hoechst (Nuclei, blue), Concanavalin A (ER, red), phalloidin & WGA (F-actin/plasma membrane, green). Scale bar indicates 50um. E. Synergy-Sensitivity scatter plot of global synergy (across each entire matrix) vs combined sensitivity score (CSS) of 36 MEKi/FAKi combinations (1296 dose variations) across 14 different cell lines. Sizing by CSS. Some combinations failed QC and were removed. Shading indicates areas of synergy (green), additivity (grey), antagonism (red). Synergy (>5 synergy score) is calculated across 4 models (Bliss, Loewe, ZIP, HSA) for concurrence.

Trametinib, GDC-0623, VS4728 and another FAK inhibitor under clinical development, AMP-945 [45] were combined as [FAKi x MEKi] variations and tested across the 14 human GSC lines, in triplicate 7x7 matrices.(Figure 5E). The data suggests that, while combined targeting of FAK and MEK with small molecules can produce spikes of strong synergy (e.g. E17 cells, Figure 5 A, B, C, D), in 29% of GBM stem cell lines (4/14) tested in this study the general trend suggest potent additivity (7/14). Synergy-Additivity-Antagonism zones in Figure 5E (Y-axis, green, grey, red shading) across 4 synergy models (Bliss, ZIP, HSA, Loewe) are indicated, versus Combined Sensitivity Score (X-axis, CSS and datapoints sized by CSS value, >30 required) which effectively quantifies synergy with potency. Across 14 glioma stem cell lines and 4 synergy models, the synergy vs CSS scores indicate a range of ‘good-to-potent’ additivity using a FAK+MEK inhibitor combination, with synergy observed in some GBM stem cells lines.

### FAK and MEK inhibitor combinations inhibit 3D GBM spheroid growth and invasion

Previous studies have shown that FAK and MEK inhibitors can both inhibit and promote invasion of GBM cell respectively [46, 47]. Therefore, we examined the effect of MEK and FAK inhibitor combinations in matrigel supported, 3D spheroid invasion assays *via* live cell imaging (Figure 6). Representative images of spheroids at 72 hrs vs 264 hrs are shown (Figure 6A), with quantification of spheroid invasion area (not spheroid size) showing trametinib alone causes an initial increase in spheroid invasion area due to a more invasive phenotype as has been reported by other studies [47]. However, time-lapse movies demonstrate this outgrowth regresses and collapses after ∼6 days exposure to compound, (Figure 6B, green data points) (Supplementary videos 1-4). Similarly, the trametinib-VS4718 drug combination shows an initial increase in spheroid invasion area, which steadily declines after 72 hrs exposure to drug (Figure 6B, red data points), and spheroid core size shows little growth overall (Figure 6A, phase images: compare far right ‘combination’ panels at 72 vs 264 hrs), indicating growth arrest. Overall, our data clearly indicates low dose targeting of MEK and FAK inhibitors in combination prevent 3D spheroid growth and invasion of human GBM models.

**Figure 6.**
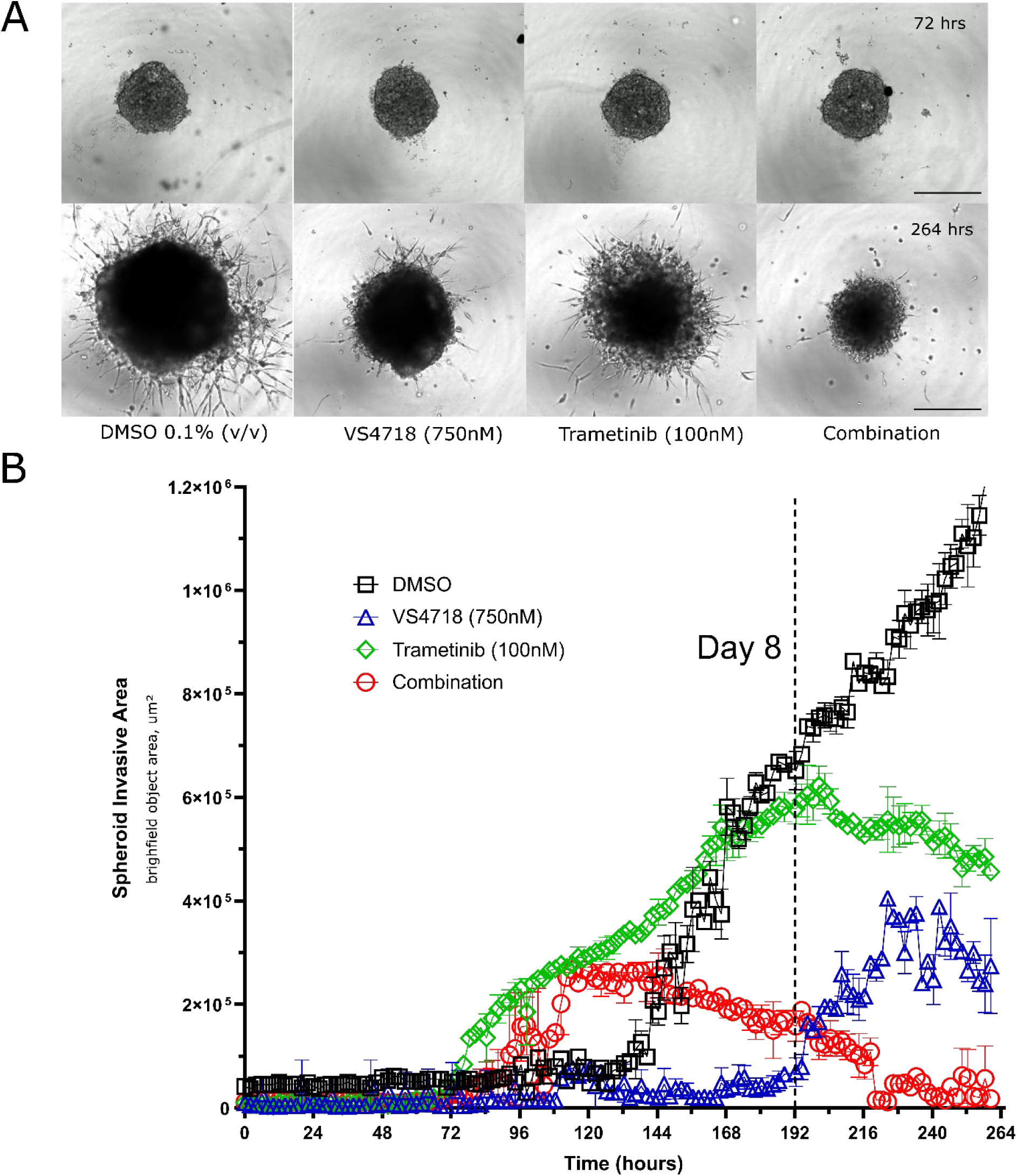
3D spheroid invasion assay with FAK and MEK inhibitor combinations. A. Representative images of live cell imaging of spheroids/invasive outgrowth at 72 (‘day 0’) and 264 hrs (∼8 days +drug) after exposure to compounds (indicated concentrations) versus DMSO control (0.1% (v/v)). Scale bar represents 500μm. B. Quantification of spheroid invasive area (exclusive of spheroid core area) over time (11 days total).

### FAK+MEK inhibitor combination profiling by Reverse Phase Protein Array (RPPA)

In order to establish tumour signalling pathway effects of the FAK+MEK inhibitor combination, beyond direct targets such as p-ERK, we profiled cell lines either resistant or sensitive to both monotherapies (E13 and E57 cells, respectively), by RPPA (127 verified antibodies, relating to common anti-cancer and adhesion signalling pathways), after treatment with either DMSO 0.1%, trametinib (100nM), VS4718 (300nM) or in combination for 24 hrs. A corresponding experiment was carried out in 3D, using the G7 GBM cells. Sample preparation and RPPA was carried out as described in methods, differential protein expression analysis across each condition was then carried out, relative to DMSO controls, quantified as Log2[FOLD] changes (Figure 7A, B).

**Figure 7.**
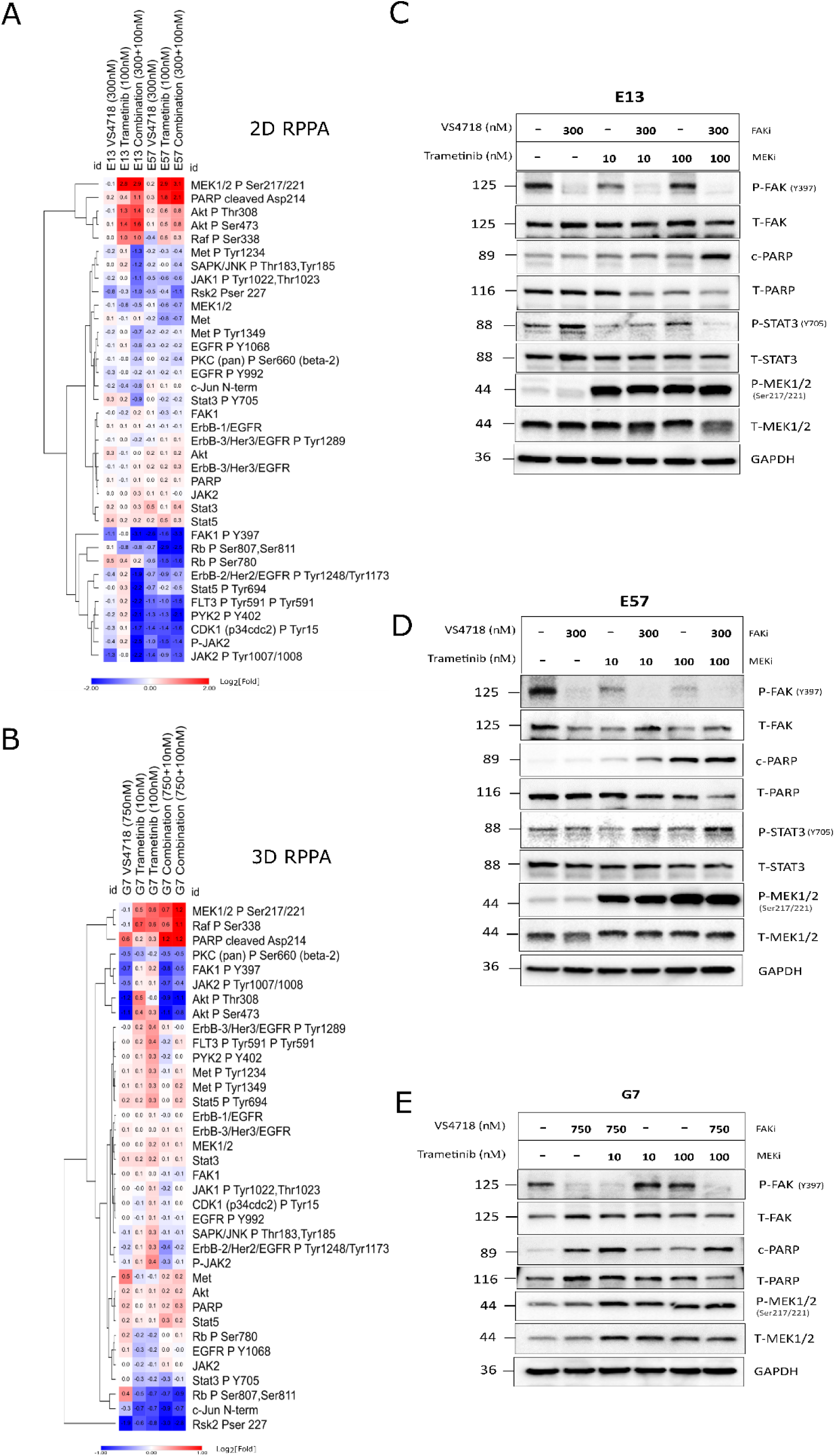
Pathway profiling of FAK+MEK inhibitor combination by Reverse Phase Protein Array (RPPA) in 2D and 3D. A.2D RPPA heatmap representation of protein expression (Log2-fold changes) in resistant (E13) vs sensitivity (E57). B. 3D RPPA heatmap representation of protein expression (Log2-fold changes) in G7 spheroids. C – E. Western blot validation of RPPA observations, including enhanced PARP cleavage with FAKi+MEK inhibitor combination,

In E13, E57 and G7 cells phosphorylated-MEK1/2 (pSer^217/224^) increased, relative to control, after trametinib alone (100nM) and in combination with VS4718 at 24 hrs (Figure 7A, top row in red plus: pMEK panel). Western blot analysis confirmed increased phosphorylation of MEK in trametinib treated cells, while pERK was still being efficiently suppressed (Figure 7C, D, E). This was evident after> 6h treatment (Supplementary Figure 5). In monotherapy resistant E13 cells, the combination shows enhanced suppression of pPYK2^Y402^, pErbB/EGFR^Y1248/1173^, pSTAT5^Y694^, pFLT3^Y591^, pRSK2^S227^ and pJAK2^Y1007/1008^. In monotherapy sensitive cells, these markers are generally already inhibited, and the combination increases this ∼2 fold. Similarly, in G7 cells under 3D culture, there were not many selective fold changes due to the combination alone, with the exception of reduced pRSK2^S227^, which is the only common factor significantly reduced in the combination arm across all 3 cell lines (Figure 7A, B), alongside increased cleaved PARP/apoptosis. The enhanced induction of apoptosis by the FAK+MEK combination was also observed across this panel of cells by western blot (Figure 7C, D, E, panel ‘c-PARP’) and live cell imaging of G7 spheroids with caspase-3 detection reagent (Supplementary Figure 6).

### *In vivo* activity

We further assessed the efficacy of the FAK+MEK inhibitor combination in an orthotopic xenograft model established using human G7 glioblastoma cells implanted intracranially in CD1 nude mice. Following tumour formation, the mice were randomized into four groups and treated with either VS4718, trametinib, their combination, or vehicle control (HPMC) for one week, followed by culling (Figure 8A). Paraffin-embedded brain sections were stained with haematoxylin and eosin and a human mitochondrial antibody to visualize and quantify the tumour area. The immunohistochemical images revealed a significant reduction in tumour area in the combination treatment group compared to either monotherapy (Figure 8B). Some positive staining was observed in non-tumour regions of the brain, which was later confirmed as non-specific binding of the antibody (supplementary figure 7). Quantitative analysis confirmed that the combination of FAK and MEK inhibitors resulted in a statistically significant reduction in tumour size relative to monotherapy arms, underscoring the potential of this combination therapy in effectively targeting glioblastoma (Figure 8C). However, it should be noted that weight loss likely due to systemic toxicity was observed in some mice within the combination treatment group by the end of the one-week treatment period (supplementary figure 4), similar to the findings in the NPE-FAK-WT model.

**Figure 8.**
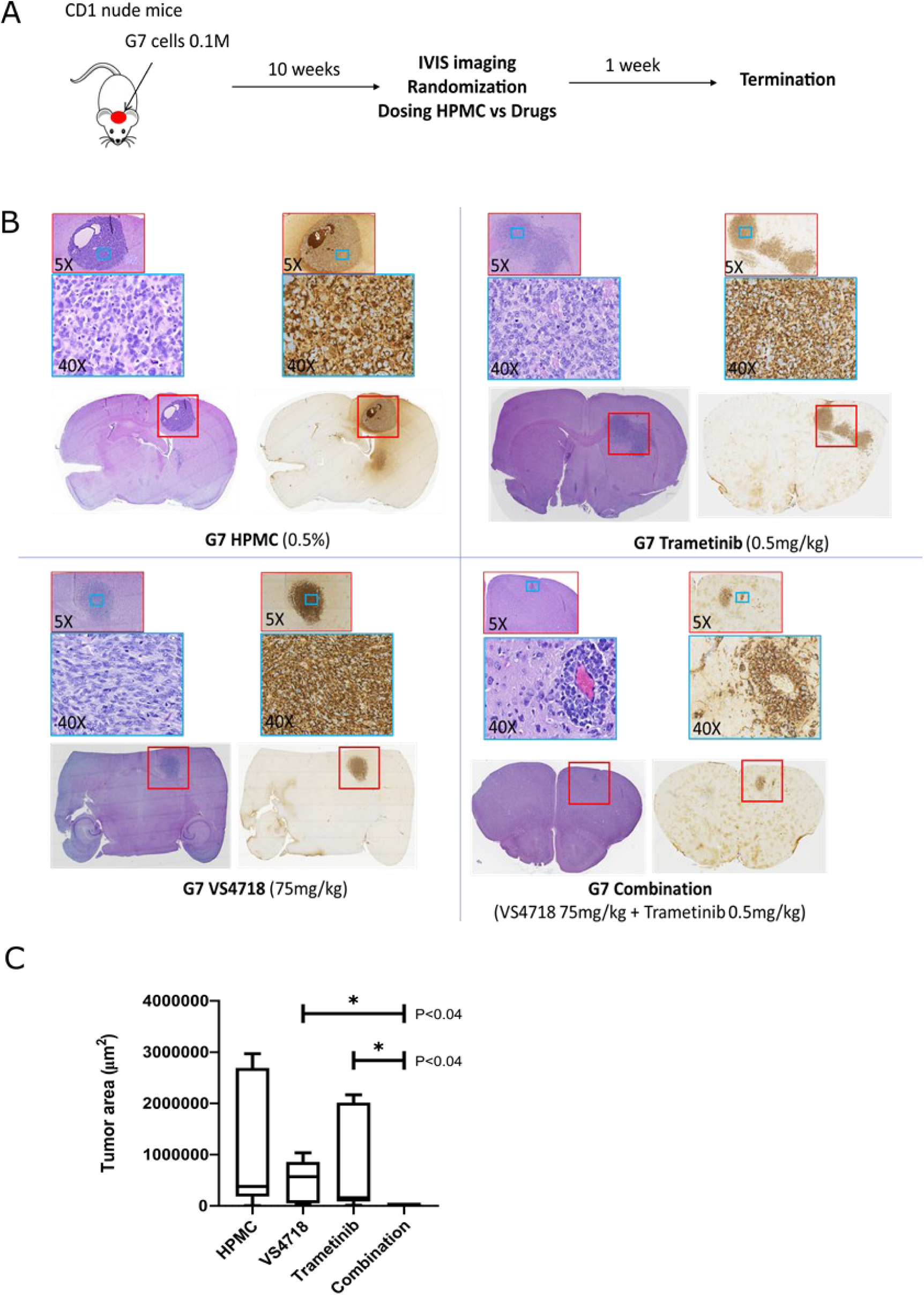
*In vivo* validation of FAK+MEK inhibitor synergy in G7 tumour bearing mice. (A) Schematic showing experimental protocol where mice with established tumours were treated with drugs (VS4718 & trametinib) and vehicle (HPMC). (B) Representative images of paraffin embedded brain sections containing G7 derived tumours stained with Haematoxylin and eosin or following immunohistochemistry with anti-human mitochondrial antibody. (C) Quantification of Tumour Area based on human mitochondrial protein staining (n=5, each group). Error bars represent the standard error of the mean (SEM). * indicates statistical significance at P < 0.04.

## Discussion

The objective of our study was to address the unmet clinical need in GBM treatment focussing specifically on discovering drug combinations that target key effectors in cellular adhesion, invasion, and migration signalling. FAK, a well characterised kinase involved in ECM remodelling, has been identified as a therapeutic target in various cancers, including GBM [18-22]. We employed an unbiased chemogenomic screening strategy in our GBM stem cell model system, using isogenic ‘kinase-deficient’ FAK cells (FAK-KD) vs wild type (FAK-WT) derived from previously established NPE parental cells (NF1^del^, PTEN^del^, EGFRvIII)[48]. We successfully screened 4 drug repurposing libraries (2586 compounds) across 4 concentrations (1000 – 30nM) yielding a 10,344 paired dataset. Through cell viability endpoints, we identified several inhibitors that demonstrated greater inhibition of cell survival in FAK-KD cells, compared to NPE-FAK-WT cells. Overlapping target mechanisms of multiple inhibitors with increased sensitivity in FAK-KD cells included HSP90 [41], MEK/MAP2K and topoisomerase-I/II inhibitors. Notably we identified two distinct MEK inhibitors, trametinib and GDC0623, that showed significantly increased sensitivity in FAK-KD cells and became the focus of our subsequent studies (Figure 1).

Our chemogenomic screening suggested that catalytic inhibitors of FAK may phenocopy FAK-KD cells and ‘synergize’ with MEK inhibitors. FAK and MEK inhibitor combinations have previously been reported in uveal melanoma [43, 44], RAS-dependent tumours [49] and NF1-related peripheral nerve sheath tumours [50], although clinical efficacy was not observed in [non-stratified] patients with advanced solid tumours, such as mesothelioma and pancreatic ductal adenocarcinoma [51, 52]. Our data in the NPE model suggested that targeting FAK and MEK against a NF1^del^ or EGFRvIII GBM background, could act synergistically. We validated this hypothesis by dose-ratio matrix testing of several FAK and MEK inhibitor combinations across multiple 2D and 3D in vitro GBM models. Synergy scores and combination indices <1 confirmed that synergistic activity is mediated through co-targeting of FAK and MEK (Figure 2A, B). Importantly, we observed additive and synergistic activity at nM concentrations of FAK and MEK inhibitors that are clinically relevant. We further validated the FAK+MEK inhibitor combination *in vivo* initially using the murine NPE intracranial transplant model (Figure 3), which showed remarkable anti-tumour efficacy resulting in substantial tumour regression. We also validated the FAK+MEK inhibitor combination using the human GBM G7 cell line that has previously been shown to maintain markers of GBM stem cells and display a growth pattern in CD1 nude mice xenografts characteristic of high-grade gliomas in patients [62] (Figure 8). Established G7 GBM tumours did partially respond to monotherapies in vivo, however, in the VS4718+trametinib combination arm, a clear and significant reduction in tumour volume was observed (Figure 8C). Although some systemic toxicity was observed with the combined use of VS4718 and trametinib the observed synergy suggests dose-sparing potential, supporting combined targeting of MEK and FAK in GBM as a valid therapeutic approach.

However, the major challenge in GBM is demonstrating efficacy across heterogeneous tumours. We profiled the FAK inhibitor, VS4718, and MEK inhibitors, GDC-0623 and trametinib, *via* dose response across a panel of 14 patient derived GBM stem cell models representing broad GBM heterogeneity (Figure 4A). Basic cell survival assays indicated unremarkable potencies as monotherapy, with a spectrum of sensitivity upon treatment with GDC-0623 and trametinib, the latter showing the strongest effects (IC50‘s ranging from 80-6000 nM) on cells carrying NF1-deletions or EGFRvIII (Figure 4B and supplementary table 1). Despite this apparent lack of potency, cellular imaging revealed distinct morphological phenotypes upon treatment with FAK and MEK inhibitor monotherapies (representative images in Figure 4C) suggesting distinct mechanisms-of-action and potential for combinatorial synthetic lethality to occur. In various [FAKi x MEKi] combinations across our glioma stem cell panel, we quantified cell survival and synergy, describing potency as a ‘combined sensitivity score’ (CSS) vs Synergy-Additivity-Antagonism scores (Figure 5E). Overall, a FAK+MEK inhibitor combination showed either synergy or potent additive effects across multiple GBM stem cell lines in 2D, and we were able to replicate these drug combination results *via* live cell imaging in a 3D spheroid invasion assay (Figure 6). Interestingly, in the 3D spheroid invasion assay, trametinib initially appeared to enhance invasion (up to 5 days of drug exposure, day 8), consistent with previous studies that raise concerns trametinib may promote invasion through strengthening of ECM adhesion contacts [47]. Nevertheless, in a 3D spheroid-Caspase-3 assay, only the FAK+MEK inhibitor combination was able to induce apoptosis over 8 days (Supplementary Figure 6) suggesting anti-tumour cytotoxic activity following long term treatment with the FAK+MEK combination overcomes any adverse activity of trametinib induced invasion.

In order to characterise signalling pathway profiles in response to the FAK+MEK inhibitor combination, we assessed several cell lines by RPPA in 2D and 3D conditions (Figure 7). A panel of well characterised antibodies (136 targets, including controls) was assayed against monotherapy sensitive and resistant cells. Interestingly, in 2D and 3D, the combination appeared to reduce pFAK^Y397^ further (relative to drug alone) and increased pMEK1/2^S217/221^, an apparent feedback signal which was confirmed by time course studies, but with continued suppression of pERK (Supplementary Figure 5). The only common factor within the drug combination arms on all 3 cell models tested here was reduced pRSK2^S227^ and enhanced apoptosis as demonstrated by western blot (cleaved PARP) (Figure 7A, B) and live capase-3 assays (Supplementary Figure 6). RSK2, a serine/threonine kinase downstream signalling mediator in the RAS/MEK/ERK pathway, has been identified as driving invasiveness/migration in glioma and other cancers [53-55], hence is a favourable downstream target and potential biomarker of the FAK+MEK inhibitor combination.

Crosstalk and compensatory signalling between the RAS/RAF/MEK and FAK/SRC signalling axis has been reported in multiple cancer cell line and preclinical tumour models [56, 57]. FAK has been shown to be activated following pharmacological inhibition of the RAS/RAF/MEK pathway in several pre-clinical tumour models and in patient tumour samples [58-60]. FAK signalling can be negatively regulated by the RAS/RAF/MEK pathway through multiple mechanisms for example, ERK-mediated phosphorylation of FAK on S910 results in FAK inactivation by dephosphorylation and focal adhesion turnover during migration. In addition,

ERK-mediated phosphorylation of calpain2 on Ser50 promote proteolysis of FAK and focal adhesion disassembly [61, 62]. In our study we observed an increase in the levels of autophosphorylated FAK-Y397 in G7 cells following treatment with trametinib in 3D in vitro culture (Fig 7B and E). However, this compensatory upregulation of FAK activity was not observed in the other GBM stem cell lines (E57 and E13) which also exhibit enhanced response to the FAK+MEK combination treatment. Rather in these cell lines the FAK+MEK combination results in enhanced downregulation of FAK-Y397 compared to FAK inhibitor alone suggesting that the MEK inhibitors are cooperating with the FAK inhibitor to repress upstream pathways promoting focal adhesion signalling.

Importantly, potent additive or synergistic activity of the trametinib + FAK inhibitor combination is observed across multiple murine and human models and is independent of BRAF^V600E^ mutation status, or specific compensatory signalling mechanisms, suggesting that this combination may serve a wider population of GBM patients than current personalised treatment propositions, such as the trametinib & dabrafenib combination, which is currently restricted to 1-2% of GBM patients that express the BRAF^V600E^ mutation [14, 15].

Combining FAK and RAF/MEK inhibitors has shown clinical activity in low grade serous ovarian cancer regardless of KRAS mutant status [64] with an ongoing registration-directed study of VS-6766 (RAF/MEK inhibitor) ± defactinib (FAK inhibitor) in patients with recurrent LGSOC (ENGOT-ov60/NCRI/GOG-3052; NCT04625270) underway [65]. These studies indicate that the FAK+MEK inhibitor combination may potentially be re-purposable for other tumours, however, improving physicochemical properties of each individual compound component, such as CNS penetrance, may be required to achieve optimal activity in GBM. Improvements in medicinal chemistry, particularly those aimed at improving the CNS penetrance of trametinib, a PgP substrate, with poor BBB permeability [66] are underway. Novel, BBB permeable MEK and ERK inhibitors for BRAF-mutant brain metastases are also being developed [67, 68]. Systemic toxicity and drug delivery across the BBB could also be overcome by emerging drug delivery devices, such as nanoparticles designed for selective delivery to tumour sites, including GBM [69, 70], Using the same NPE glioma model used in the current studies we have recently reported on another integrin adhesome node, namely integrin linked kinase (ILK), which controls transcriptional plasticity [71]. Depletion of ILK drives conversion of GBM stem cells from a heterogenous population to a more homogenous one, thus the FAK+MEK combination treatment may also target plasticity to provide broad activity across heterogenous GBM stem cell models. Future studies to promote translation of FAK+MEK inhibitor combination therapy into GBM clinical trials include broader profiling across a larger panel of GBM stem cell models and correlation of phenotypic outcome with gene expression profiles to identify molecular biomarkers that predict optimal synergy to support patient stratification for personalised medicine strategies. Identification of proof-of-mechanism and target engagement biomarkers to guide optimal dosing and scheduling of the combination and preclinical toxicology, considering targeted drug delivery strategies, if necessary, to decipher the therapeutic window and safety margins of FAK+MEK inhibitor combination will also build confidence in the clinical potential of FAK+MEK inhibitor combination therapy in GBM.

## Acknowledgements

This work was funded by a joint Cancer Research UK (C42454/A28596) and The Brain Tumour Charity award (GN-000676) to D.E., N.O.C. and M.C.F. G.M.M and the Glioma Cellular Genetics Resource were supported by the Cancer Research UK (CRUK) Centre Accelerator Award (A21922).

