## Supplementary Figures for "Drug combinations targeting FAK and MEK overcomes tumour heterogeneity in glioblastoma"

### Supplementary Figure 1.

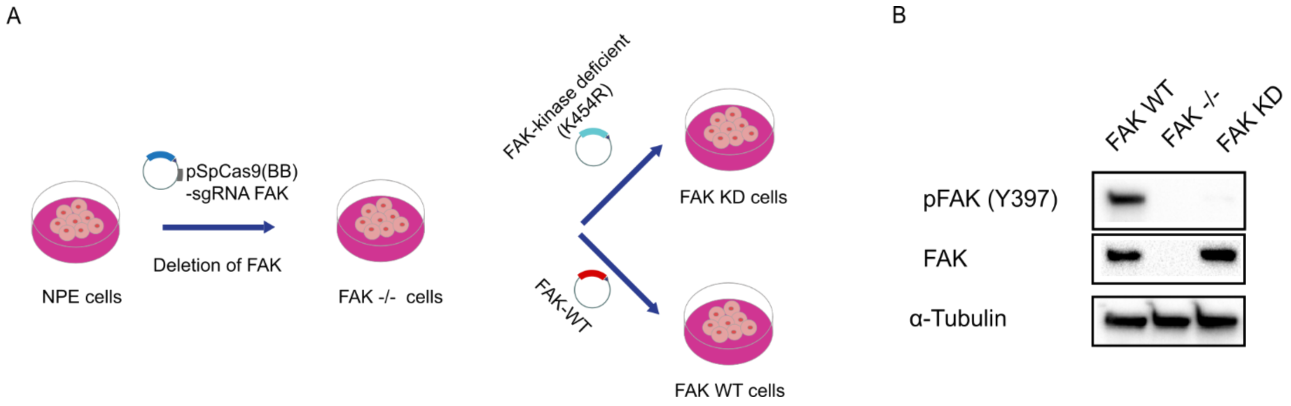


**Supplementary Figure 1:** Generation of FAK kinase deficient NPE cells. A. Schematic of strategy used to generate FAK_kinase deficient (K454R) cells (FAK KD) from NPE cells. B. Immunoblot of FAK WT, FAK -/- and FAK KD cells showing pFAK (Y397), and FAK expression. α-Tubulin was used as a loading control.

### Supplementary Fig 2:

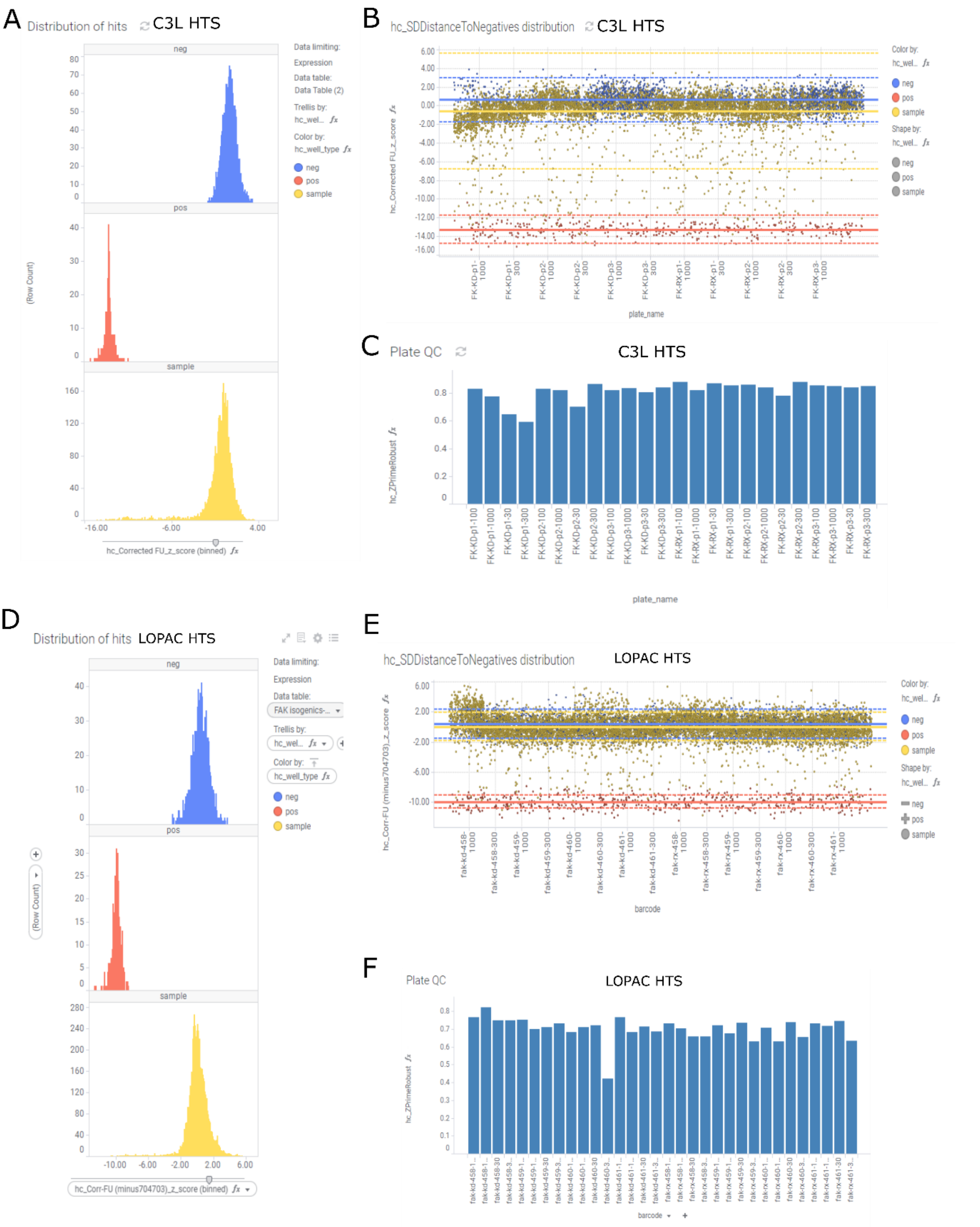


**Supplementary Figure 2**: Representative data of normalised high throughput screening analysis. A. C3L Library Normalised frequency distribution (Z-score) of negative controls (blue, 0.1% DMSO), positive controls (red, 1uM staurosporine) and samples (yellow). B. C3L Scatter plot representation of normalised data (Z-score) across each plate. Distribution lines (solid) represent median z-score with dispersal range (dotted lines; 3 MAD 'median-absolute deviation') for negative controls (blue, 0.1% DMSO), positive controls (red, 1uM staurosporine) and samples (yellow). C . C3L Z-prime data across each plate, ideally >0.5 for excellent assay quality. D. LOPAC Library Normalised frequency distribution (Z-score) of negative controls (blue, 0.1% DMSO), positive controls (red, 1uM staurosporine) and samples (yellow). E. LOPAC Scatter plot representation of normalised data (Z-score) across each plate. Distribution lines (solid) represent median z-score with dispersal range (dotted lines; 3 MAD 'median-absolute deviation') for negative controls (blue, 0.1% DMSO), positive controls (red, 1uM staurosporine) and samples (yellow). F . LOPAC Z-prime data across each plate, ideally >0.5 for excellent assay quality.

### Supplementary Fig 2 (cont'd):

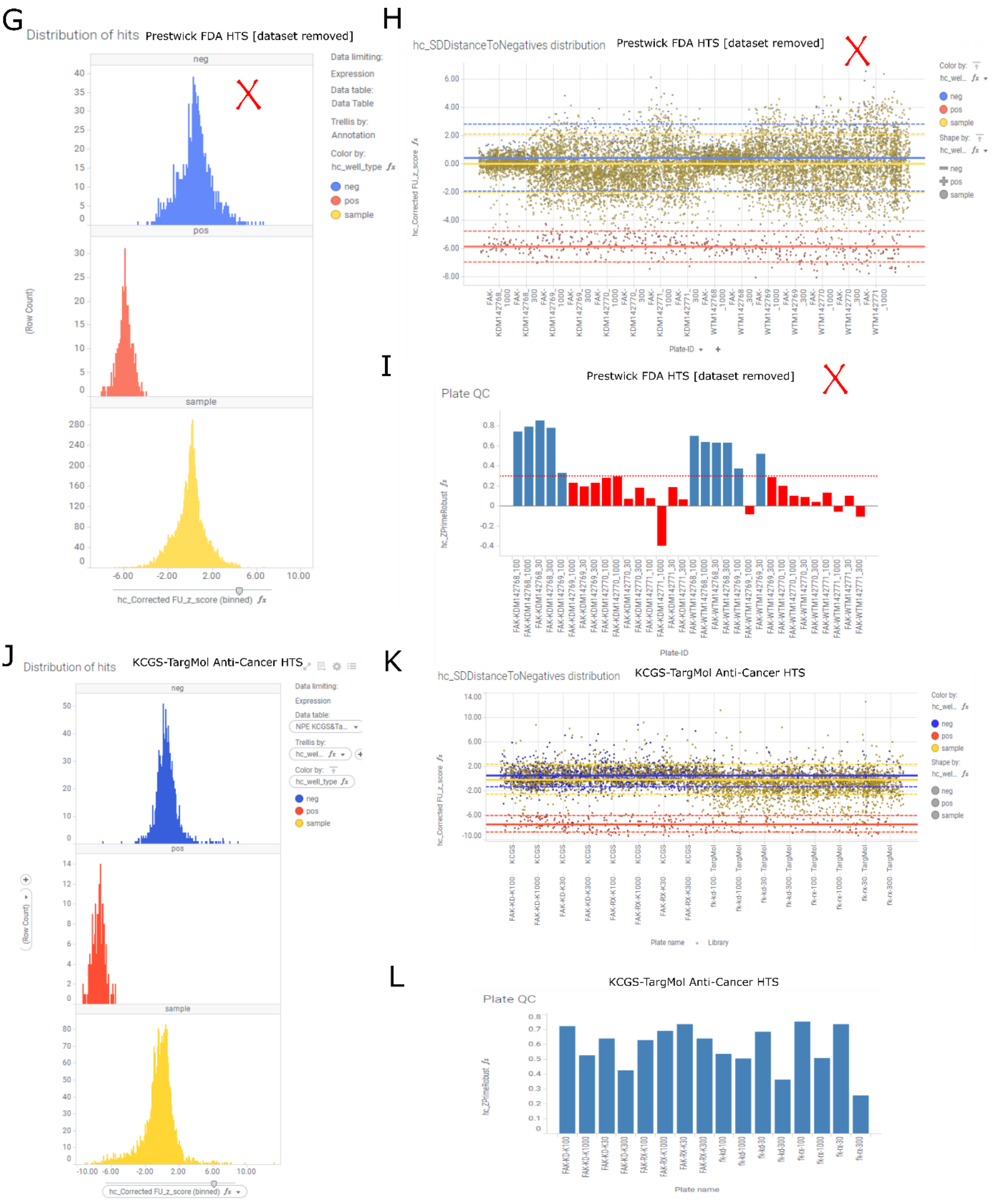


**Supplementary Figure 2 cont'd** : Representative data of normalised high throughput screening analysis. G. Failed QC. Data rejected. Prestwick FDA Library Normalised frequency distribution (Z-score) of negative controls (blue, 0.1% DMSO), positive controls (red, 1uM staurosporine) and samples (yellow). H. Failed QC. Data rejected. Prestwick FDA Scatter plot representation of normalised data (Z-score) across each plate. Distribution lines (solid) represent median z-score with dispersal range (dotted lines; 3 MAD 'median-absolute deviation') for negative controls (blue, 0.1% DMSO), positive controls (red, 1uM staurosporine) and samples (yellow). I. Failed QC. Data rejected. Prestwick FDA Z-prime data across each plate, ideally >0.5 for excellent assay quality. J. KCGS-TargMol Anti-Cancer Library Normalised frequency distribution (Z-score) of negative controls (blue, 0.1% DMSO), positive controls (red, 1uM staurosporine) and samples (yellow). K. KCGS-TargMol Anti-Cancer Library Scatter plot representation of normalised data (Z-score) across each plate. Distribution lines (solid) represent median z-score with dispersal range (dotted lines; 3 MAD 'median-absolute deviation') for negative controls (blue, 0.1% DMSO), positive controls (red, 1uM staurosporine) and samples (yellow). L . KCGS-TargMol Anti-Cancer Library Z-prime data across each plate, ideally >0.5 for excellent assay quality.

### Supplementary Fig 3:

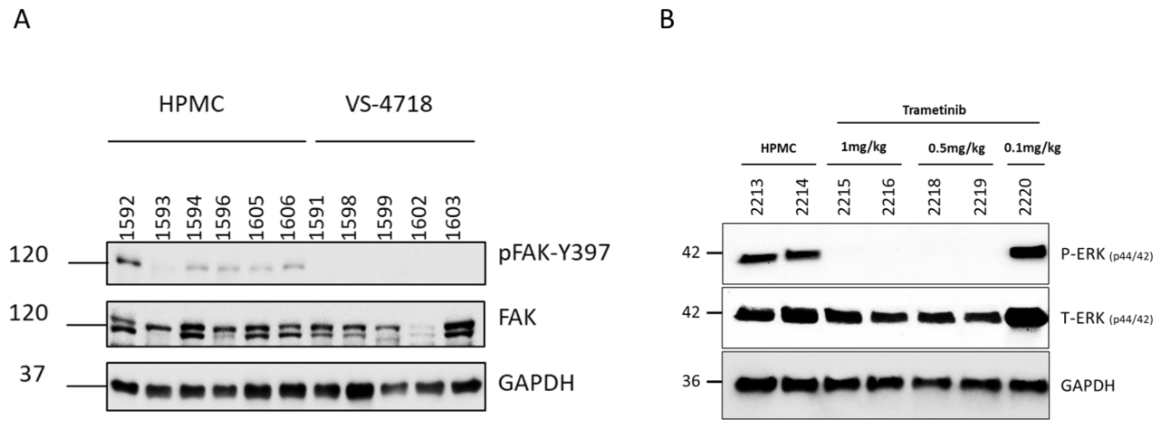


**Supplementary Figure 2:** In vivo biomarker modulation by FAK and MEK inhibitors reflect target engagement in mice brains. Western blot analysis of tumour lysates showing inhibition of (A) P-FAK in VS4718 75mg/kg treated tumours and (B) P-ERK in Trametinib treated tumours.

### Supplementary Fig 4:

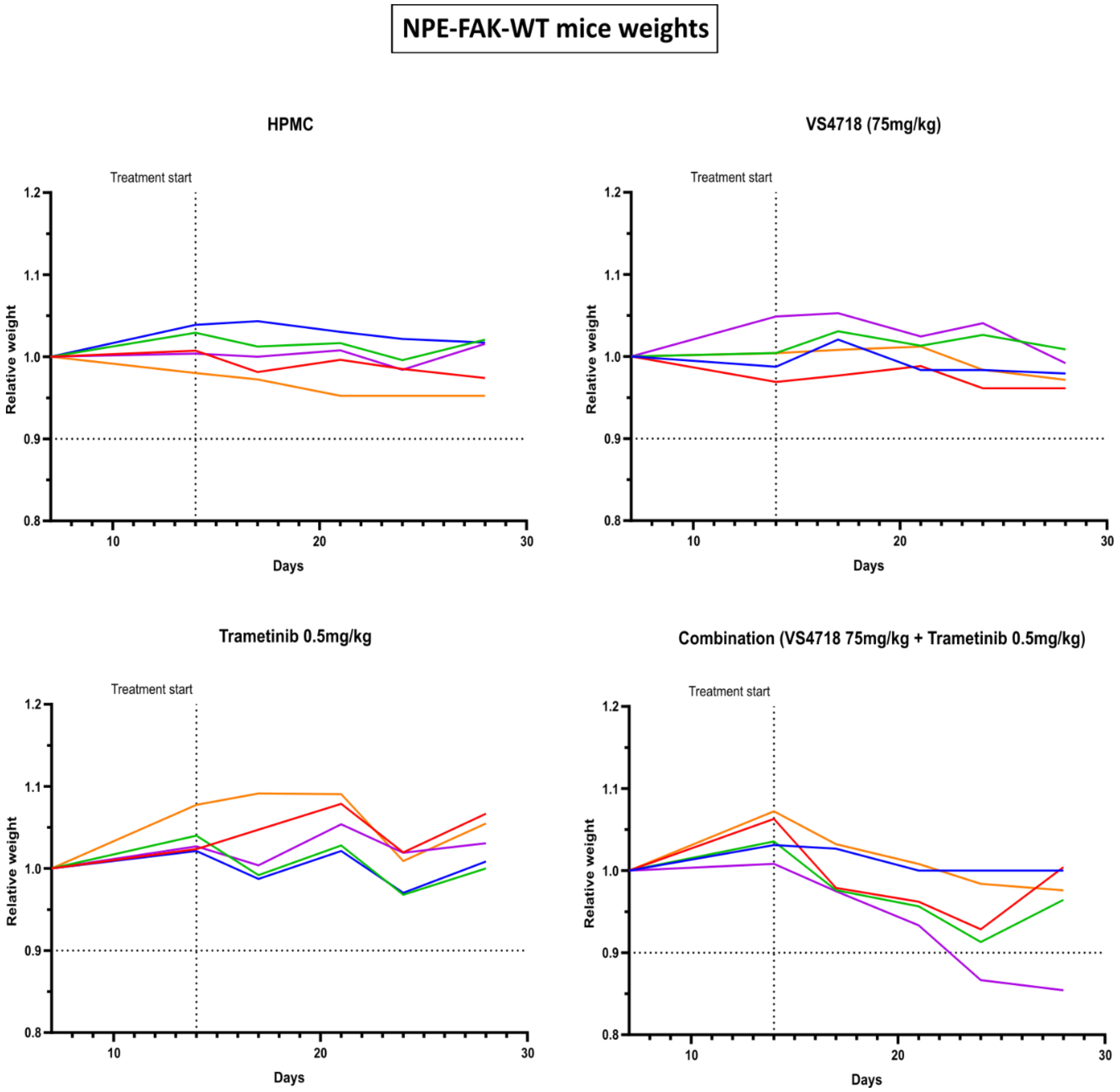

#### G7 mice weights

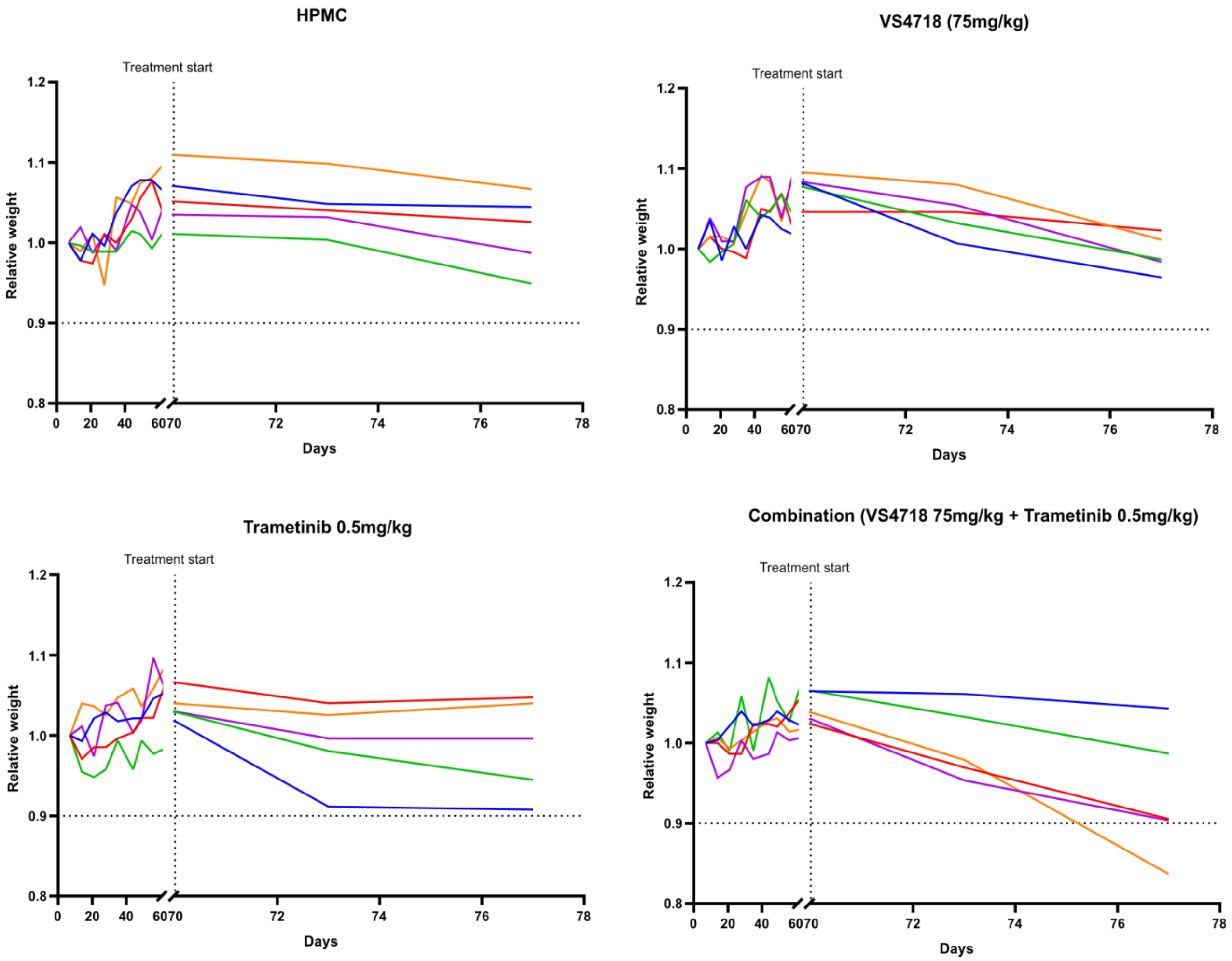


**Supplementary Figure 4:** Body weight measurements of NPE-FAK-WT and G7 tumour bearing mice treated with the indicated drugs over the course of two and one week, respectively via oral gavage. Relative weight was calculated by dividing the weight of each mouse on a given day by its weight on day 1. Each coloured line represents one mouse.

### Supplementary Fig 5:

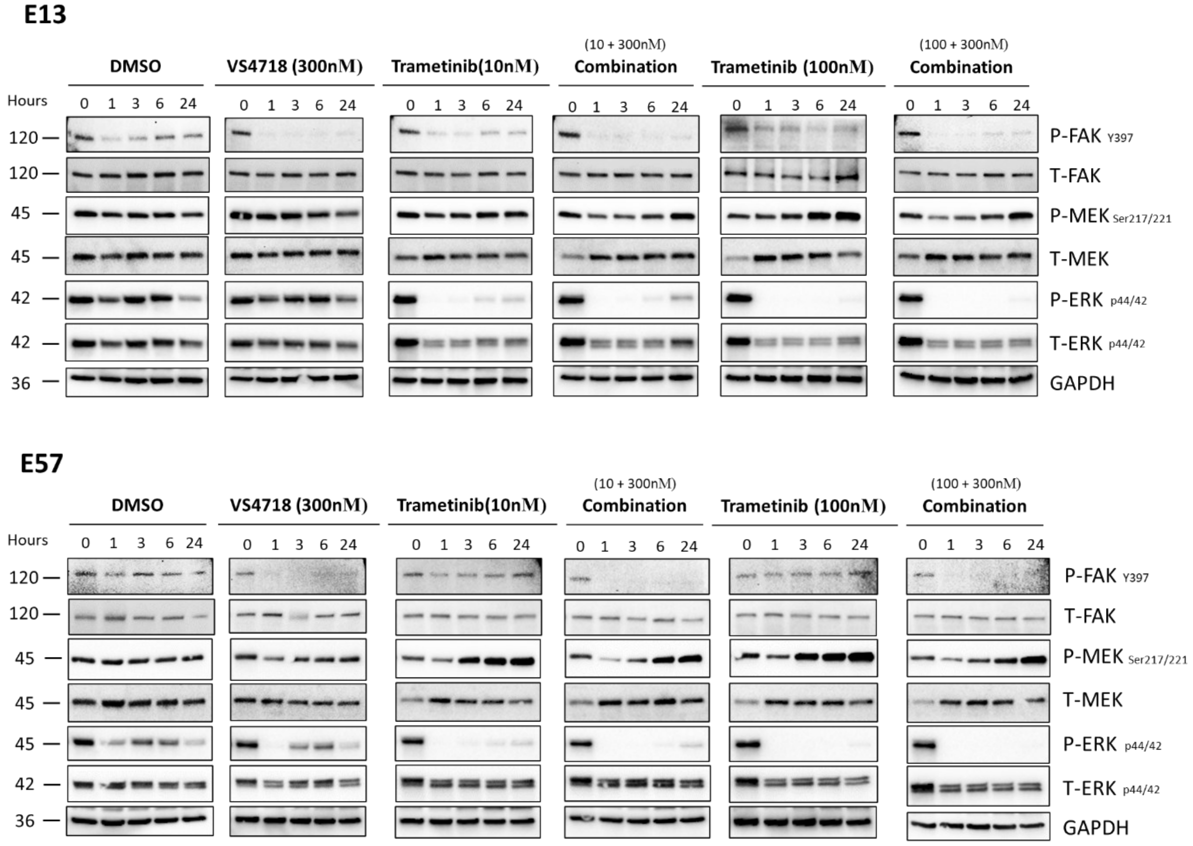


**Supplementary Figure 5:** Examining the time course of FAK, MEK and ERK Phosphorylation following treatment with indicated drugs in E13 and E57 cells

### Supplementary Figure 6.

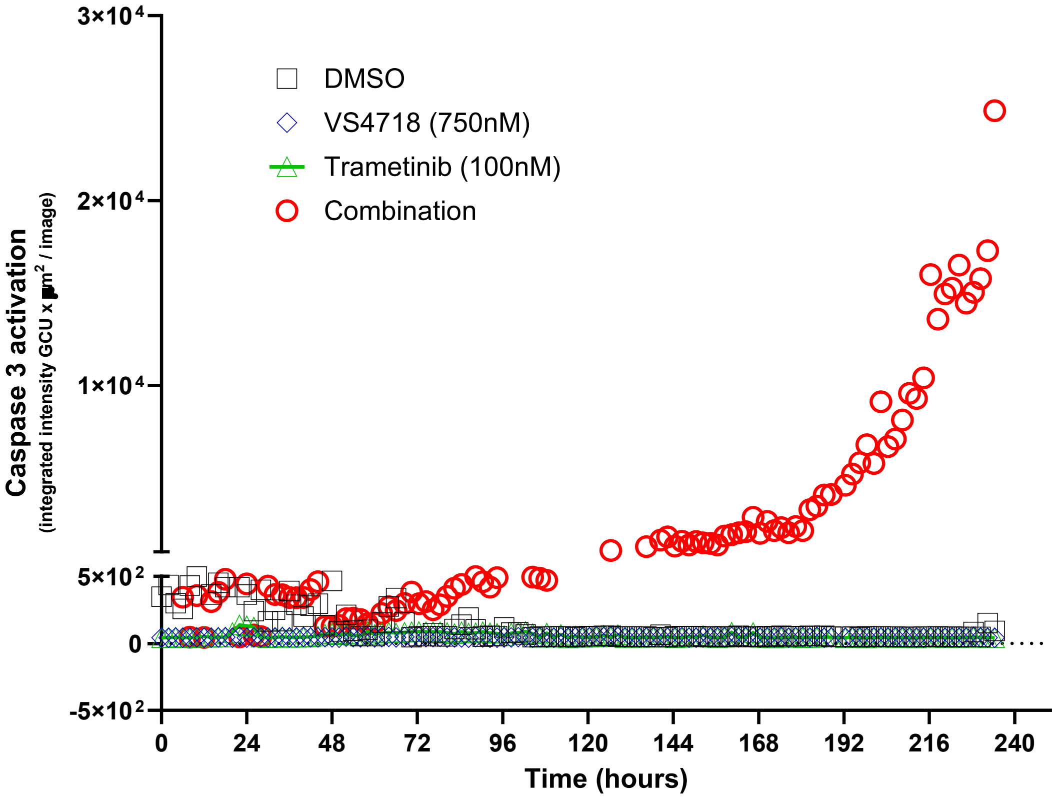


**Supplementary Figure 6:** Treatment with FAK-MEK inhibitors combination causes activation of caspase-3 in G7 GBM spheroids. Caspase-3 activation was determined using BioTrac 530 Red Caspase-3 Dye (Sigma-Aldrich) according to manufacturer's instructions.

### Supplementary Figure 7.

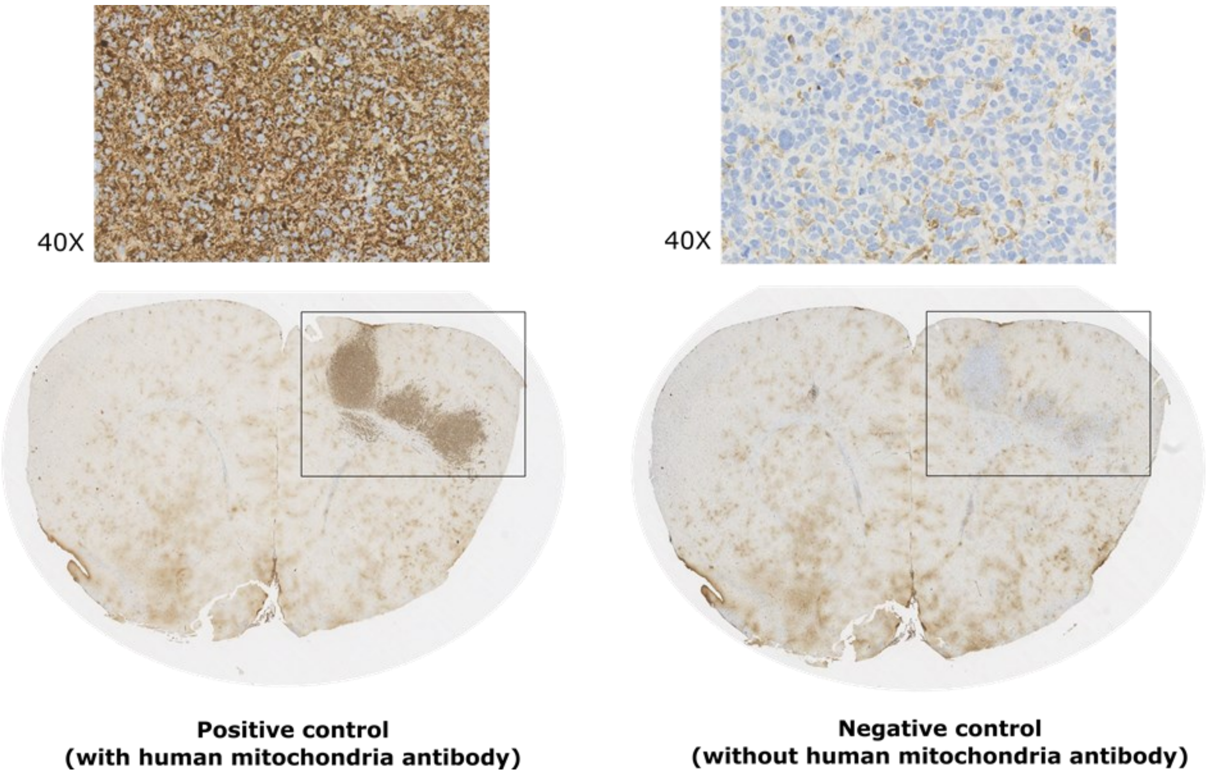


**Supplementary Figure 7:** Immunohistochemistry using an anti-mitochondria antibody with positive and negative controls to assess specificity. The positive control (left) shows strong mitochondrial staining, while the negative control (right) displays minimal non-specific binding. Boxed regions highlight staining patterns for comparison.
